# Stearic acid enhances membrane fluidization and peptidoglycan stiffness to promote stability of Gram-positive bacteria

**DOI:** 10.64898/2026.03.10.710747

**Authors:** Srividhya Parthasarathi, Sachinkumar Jayantibhai Joshi, Jaydeep K. Basu, Rakesh Vaiwala, K. Ganapathy Ayappa, Morris Waskar, Srikala Kumaran, Anindya Dasgupta

## Abstract

Saturated fatty acids such as stearic acid (SA) can exhibit both antimicrobial and growth-promoting effects on bacteria, depending on their concentration and chemical structure. However, the physical properties of the bacterial cell envelope in response to such molecules remain under-explored compared to their biochemical pathways. In this study, a comprehensive investigation is presented on the interaction of SA with the Gram-positive bacterium, *Staphylococcus epidermidis* (*S. epi*). SA alters bacterial growth, reflected in a higher maximum specific growth rate, a shorter lag phase, and an extended exponential phase, consistent with a prebiotic effect. Using fluorescence correlation spectroscopy and fluorescence lifetime imaging microscopy, we show that SA incorporation leads to significant fluidization of the lipid membrane, characterized by enhanced lateral diffusion and reduced membrane viscosity. Coarse-grained molecular dynamics (CG-MD) simulations demonstrate spontaneous insertion of SA into the membrane and a significant increase in mean-square displacement after insertion, supporting our experimental observations. Importantly, atomic force microscopy measurements show an increase in cell-envelope stiffness, reflected by a higher Young’s modulus which can be attributed to modulations in the glycan-peptide linkage density based on earlier studies that correlate stiffness changes to peptidoglycan (PG) crosslinking in Gram-positive strains [1]. These results provide direct evidence linking membrane fluidization induced by SA and increased cell wall stiffness due to transport modifications in the membrane mediated PG synthesis pathways to enhance bacterial cell viability.

## I. INTRODUCTION

Bacterial cells possess a densely packed cytoplasm enclosed by a multilayered cell envelope. In Gram-positive bacteria, the cell envelope includes an overlying thick peptidoglycan (PG) that preserves cell shape and prevents osmotic lysis [2, 3], and a phospholipid bilayer underneath that functions as a selective permeability barrier, maintaining ion gradients and facilitating the transport of nutrients and signalling molecules [4, 5]. The physical parameters of bacterial membranes play a crucial role in maintaining cellular integrity, ensuring selective permeability to molecules such as nutrients, and enabling adaptive responses essential for bacterial survival [6–9]. Importantly, the dynamical and mechanical properties of the bacterial membrane, such as lipid packing, viscosity, lateral diffusion, and rigidity, are strongly influenced by external amphiphilic molecules, including fatty acids [10–13] Fatty acids are organic compounds containing a carboxylic acid group attached to a long aliphatic chain, which can be saturated, unsaturated, or branched. They can act as antimicrobial agents [14–17] and also serve as modulatory molecules, exerting either inhibitory or beneficial effects depending on their chemical structure and concentration [18– Although unsaturated fatty acids have been extensively investigated for their antibacterial properties [23, 24], the functional roles of saturated fatty acids (SFAs) in bacterial systems remain relatively less explored. SFAs play crucial roles in maintaining bacterial membrane integrity and participate in lipid metabolism, which can influence energy balance and the stability of microbial communities [25]. Most studies on SFAs have focused on their antimicrobial properties. For example, in one study, SFAs were encapsulated in liposome carriers, demonstrating bactericidal activity against multidrug-resistant *Staphylococcus epidermidis* (*S. epi*) and vancomycin-resistant *Enterococcus faecalis*[26]. In another experimental study, stearic acid (SA), a type of SFA, formed nanostructured arrays upon recrystallization on the surface of highly ordered pyrolytic graphite, and these arrays exhibited antimicrobial activity against *Pseudomonas aeruginosa* and *Staphylococcus aureus (S*.*aureus)* [27]. In addition to these antimicrobial effects, recent research indicates that SFAs have beneficial effects on gut health, improve liver function, and shape the composition of the gut microbiota in humans [28]. However, these studies have mainly examined the interaction of SA with bacterial membranes—either as part of liposome carriers for drug delivery or as surface coatings on graphite substrates—while the direct effects of SA on bacterial cells themselves remain largely unexplored.

Due to the complex organization of the bacterial cell envelope, developing realistic models of bacterial membranes has become an important focus of current research [10, 29, 30]. Although model membranes are relatively easy to handle, studying live bacteria remains challenging due to the complex, multi-component nature of their membranes. Such studies are crucial, as they provide real-time insights into the dynamic and mechanical properties of the bacterial cell envelope. These properties play key roles in cell division, motility, adhesion, maintenance of selective permeability, and essential processes such as nutrient uptake, stress response, and membrane integrity [31–35].

In recent years, atomic force microscopy (AFM) based studies have provided important insights into antimicrobial interactions with the PG layer. In an AFM study on *S*.*aureus*, Bailey et al. [36] identified two distinct mechanical regimes: stiffness measured at low indentation depths reflects the local stiffness of the cell wall, while stiffness at higher indentation depths is dominated by whole-cell mechanical responses, primarily influenced by turgor pressure. In another AFM study on *S. epi*, mechanical parameters were extracted from AFM force-indentation data using various models, such as the Hertz model, thin elastic shell models, and composite shell–turgor pressure models. Similar to the study mentioned above, it was found that parameters obtained at low indentation depths mainly reflect the intrinsic elasticity of the bacterial cell wall, whereas measurements at larger indentation depths are controlled by whole-cell mechanics, including turgor pressure and cell geometry, in agreement with the observations reported by Bailey et al. [32]. In many cases, exposure to antibiotics or cell-wall–targeting agents is associated with a reduction in Young’s modulus, reflecting mechanical weakening of the envelope due to structural damage. For example, AFM measurements on *Escherichia coli* (*E. coli*) have shown a decrease in Young’s modulus following treatment with *β*-lactam antibiotics such as ampicillin, while studies on *S*.*aureus* exposed to lysostaphin—an enzyme that specifically cleaves peptidoglycan—report reduced cell stiffness [37, 38]. Such observations indicate that antimicrobial stress commonly leads to softening of the bacterial cell wall, consistent with degradation or disruption of the PG network. However, compared with the extensive literature on antimicrobial compounds, the direct interactions of growth-promoting molecules with the Gram-positive bacterial membrane remain relatively underexplored. In particular, a direct correlation between biophysical changes in various components of the membrane with the eventual cell viability remains sparsely explored [36, 39–42].

In this study, we investigate the role of SA as a modulator of bacterial growth and examine how it influences the structural and mechanical properties of the cell membrane in *S. epi*. Bacterial growth kinetics were measured using a plate reader, allowing continuous monitoring of optical density over time and modelled with the Gompertz framework to extract key parameters such as maximum specific growth rate, lag phase duration, and exponential growth time. To probe the physical response of the lipid membrane, we employed fluorescence correlation spectroscopy (FCS) and fluorescence lifetime imaging microscopy (FLIM), which provide complementary information on lipid mobility and membrane viscosity following SA treatment. Coarse-grained molecular dynamics (CG-MD) simulations were performed to understand the insertion energetics of SA into *S. epi* membrane and its effects on lipid diffusion. To quantify SA-induced alterations in the PG layer, Young’s modulus derived from AFM measurements was employed as a mechanical parameter. Overall, these findings indicate that SA can alter bacterial membrane dynamics, affecting both lipid fluidity and PG rigidity, and may drive prebiotic effects by supporting growth, structural integrity, and cellular stability.

## II. RESULTS

### Prebiotic effects of SA from growth kinetics measurements

Growth kinetics of *S. epi* in the presence of SA was evaluated using a 96-well plate assay, with four independent trials in each experimental run to ensure reproducibility. Data from additional sets are provided in the Supplementary Fig. S1 and S3. Optical density at 620 nm (OD_620_) was recorded at regular time intervals, and the resulting curves were plotted as OD_620_ versus time (Figure 1a). The increase in OD_620_ reflects a corresponding rise in cell density and serves as a reliable indicator of bacterial growth. Comparison of measurements before and after SA treatment indicates that SA enhances the overall growth of *S. epi*,evidenced by the higher OD values following treatment. Figure 1b shows the relative change at any time, *t* defined as the normalized variation in OD, (*OD*(*t*) − *OD*(*t*_*o*_)*/OD*(*t*_*o*_)) that quantifies the fractional change in growth rate as a function of incubation time. A distinct increase in the relative change is observed at higher concentrations of SA at the earlier time points. These findings suggest that SA may have a potential prebiotic effect on *S. epi*, defined as the selective ability of a compound or molecule to enhance the growth, survival, or metabolic activity of a specific bacterial species.

**FIG. 1.**
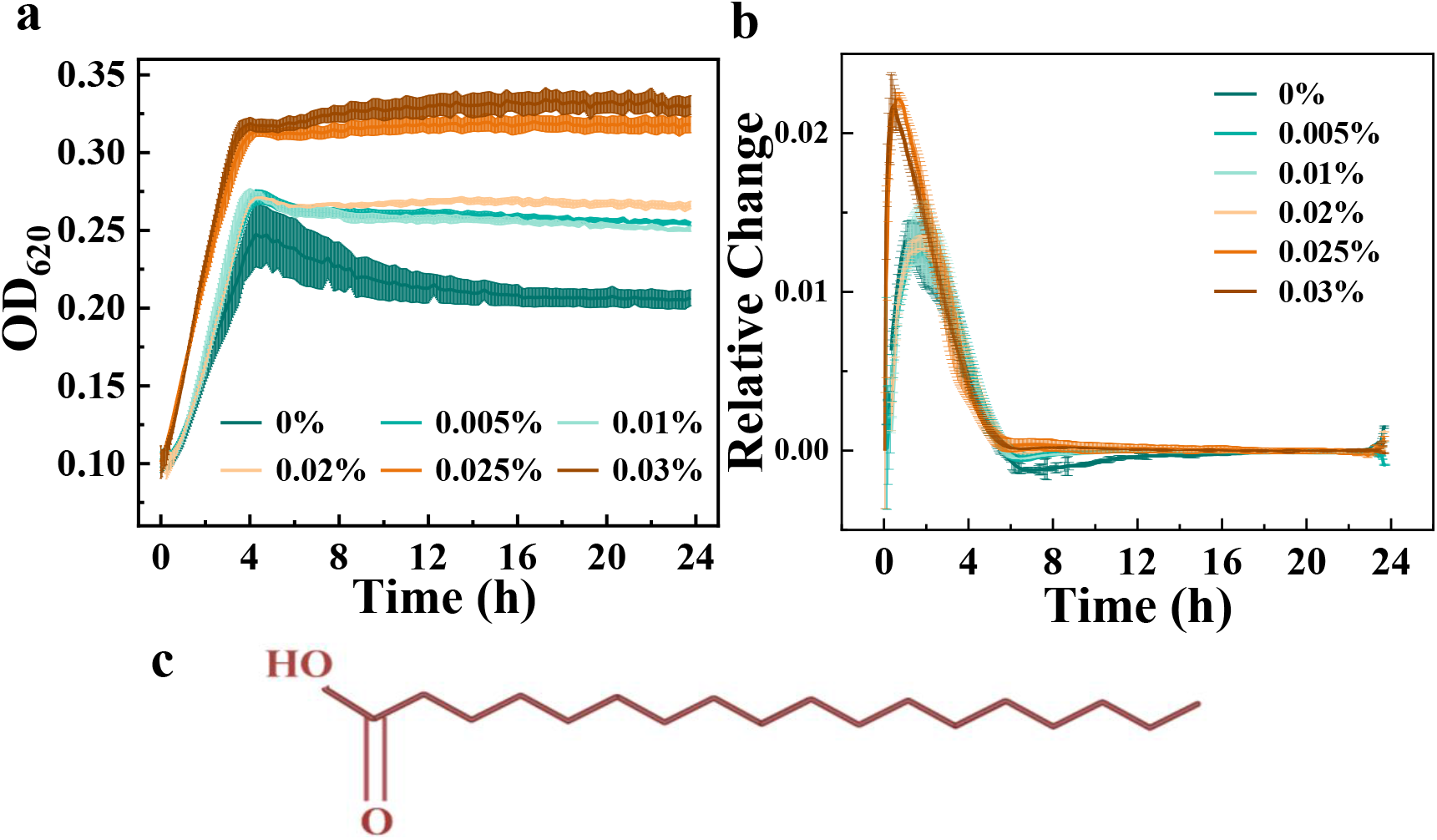
(**a**) Growth kinetics curves of *S. epi* with and without SA at various tested concentrations, showing increased optical density (OD) and a longer stationary phase upon SA treatment. (**b**) Relative growth rates extracted from the kinetics curves, highlighting the faster growth rate along with an increase in OD at higher SA concentrations. (**c**) Molecular structure of SA.

To quantitatively evaluate the effect of SA on the growth dynamics of *S. epi*, the growth kinetic data were analysed using the modified Gompertz model [43–45] given in Equation 1. This model describes microbial growth using three sigmoidal fit parameters (a, b, and c) as given below,

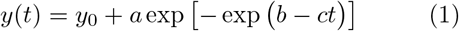

where *y*(*t*) represents the OD at time *t, c* is the growth rate and *a* is the asymptote at long time. In the original Gompertz solution, the constant *b* is related to both *a* and the initial concentration *y*_*o*_ (see Supplementary material). In the modified Gompertz model, *y*_0_ is included in Gompertz equation as an additive constant. Note that as *t* → ∞, *y* = *a* + *y*_0_. The maximum specific growth rate, *µ*_max_ = *ace*^−1^, corresponding to the fastest cell division rate during the exponential growth phase. The lag time, *λ* = (*b* − 1)*/c*, refers to the time for cells to adapt to the medium. The duration of the exponential phase is estimated using *T*_exp_ = *a/µ*_max_. These parameters serve as quantitative indicators of growth kinetics to assess the potential prebiotic effect of SA on bacteria.

We fitted Eq. 1 using the Marquardt-Levenberg algorithm in Origin with *y*_0_ used as a fourth parameter. This was found to improve the quality of the fit at short times and reliably captured the sigmoidal features of the kinetics with goodness-of-fit values ranging from *R*^2^ = 0.97 to 0.99. The model fits for SA concentrations of 0 and 0.03% are shown in Figure 2 and the fit parameters for the complete set of SA concentrations is given in Table 1 of the Supplementary material. As shown in Figure 2, the maximum specific growth rate (*µ*_max_) of *S. epi* increased with increasing concentrations of SA. An increased *µ*_max_ shows that cells divide more rapidly during the exponential phase, suggesting that SA enhances the metabolic efficiency of the bacterium under the experimental conditions. In parallel, the lag phase duration (*λ*) decreased steadily with increasing SA concentration, indicating that cells adapt more quickly to the growth medium in the presence of SA. In particular, the exponential phase duration (*T*_exp_) increased with SA supplementation, indicating that the population maintained active division for a longer duration. These trends suggest that SA promotes faster adaptation, accelerated growth, and sustained proliferation of *S. epi*, consistent with a potential prebiotic effect [46]. Additional growth descriptors derived from the Gompertz model—including inflection time (*T*_inf_), cell density at the inflection point (*OD*_inf_), generation time (*T*_*g*_) and the number of generations per hour (*k*)—are summarized in Table 1 in supplementary material with few other parameters (Supplementary Fig. S2). Importantly, the same SA concentrations induce only marginal changes in the growth kinetics of the Gram-negative bacteria *E. coli* (Supplementary Fig. S3), indicating the specificity of SA’s effects on *S. epi*.

**FIG. 2.**
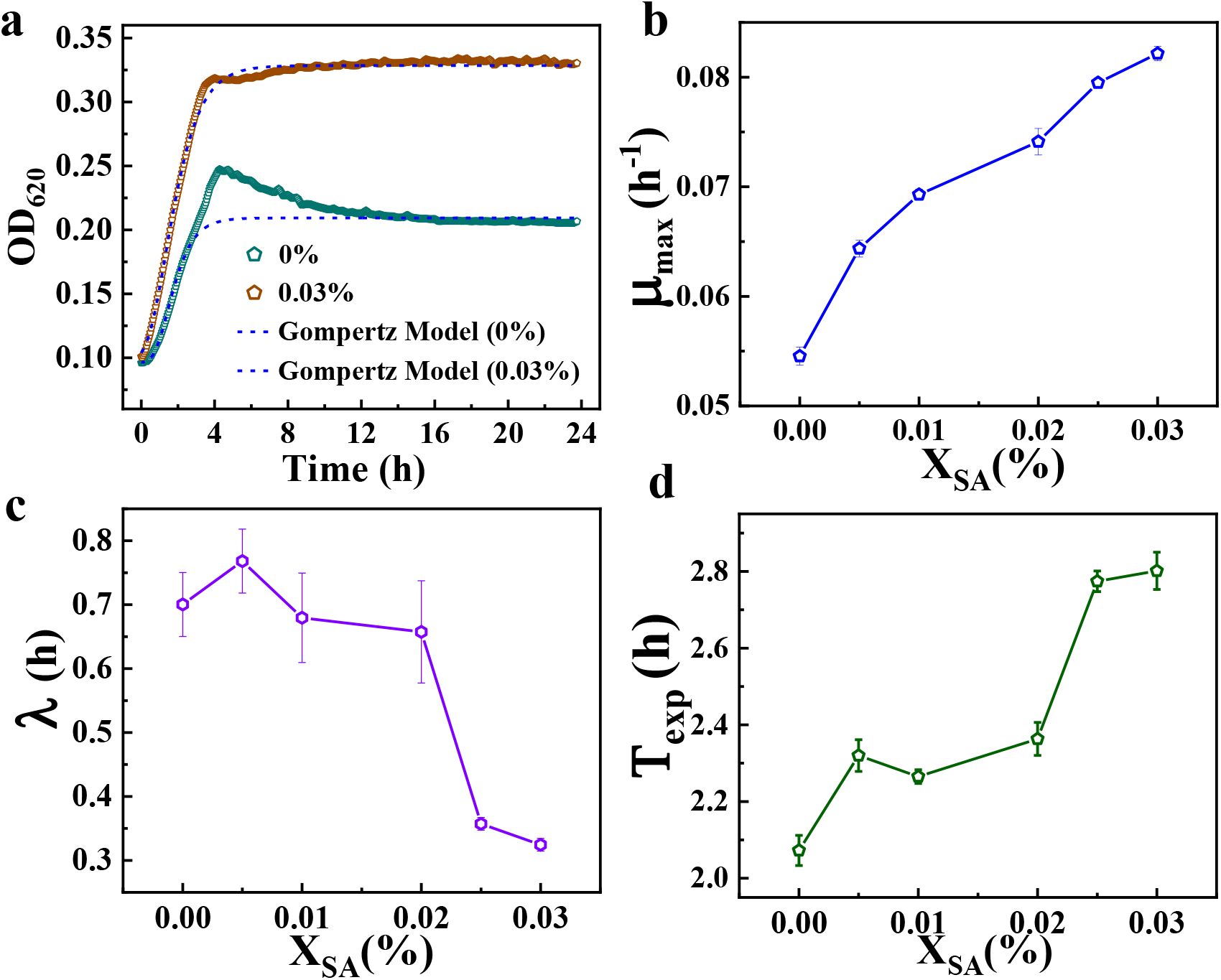
(a) Growth kinetics of *S. epi* in the absence and presence of SA. (b) Maximum specific growth rate (*µ*_max_), lag phase duration (*λ*), and (d) exponential phase duration (*T*_*exp*_) as a function of SA concentration.

### Interaction of SA with bacterial membrane: Confocal imaging and FCS

The experiments involving bacterial cells were conducted during the late logarithmic growth phase, in the presence or absence of SA, at 37^*◦*^C with continuous shaking. Nile Red (NR), a lipid-specific hydrophobic dye, was used for imaging the live bacterial membrane. Bacteria, both with and without SA treatment, were labeled with NR (Figure 3a) to investigate the effects of SA on membrane dynamics. No noticeable changes in the shape or size of the bacteria were observed after incubation with SA. Membrane dynamics were quantified using FCS on NR labeled cells. The autocorrelation profiles for bacteria with and without SA (0.03%) are presented in Figure 3b, comparing the lipid diffusion dynamics before and after SA incubation. A histogram of the diffusion coefficient (*D*) measured on the membrane is shown in Figure 3c. *D* was calculated from the autocorrelation profiles using the following equation

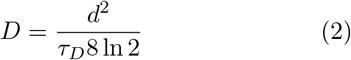

where *d* represents the point spread function of the Gaussian beam. *τ*_*D*_ is the diffusion time obtained from the autocorrelation function, from which *D* was calculated.

**FIG. 3.**
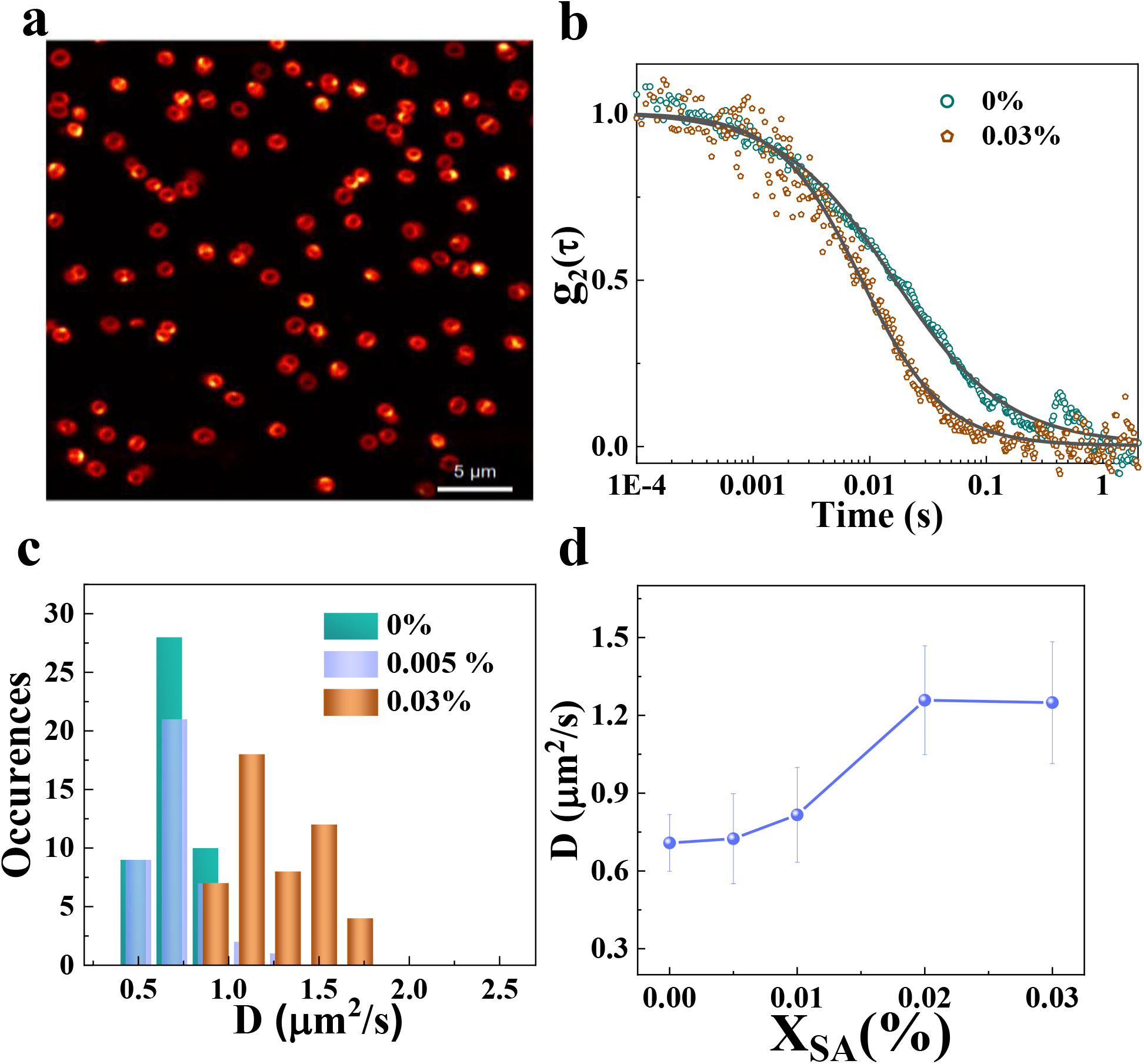
(a) *S. epi* cells labelled with Nile Red, clearly staining the bacterial membrane. (b) Autocorrelation profiles were plotted with and without SA incubation. A shift to faster dynamics is observed upon SA treatment, suggesting increased membrane fluidity. (c) Histogram of *D* at various SA concentrations, showing distribution shifts indicative of changes in membrane dynamics. (d) Concentration-dependent increase in *D* with increasing concentration of SA, suggesting increased inner membrane fluidization.

The average *D* in pristine cells was 0.71 *±* 0.12 *µ*m^2^/s, which increased to 1.25 *±* 0.24 *µ*m^2^/s after incubation with 0.03 % SA — an approximate increase of 78%, indicating membrane fluidization. Similar changes were also observed in the stationary phase; however, the absolute values were lower than those in the late log phase, possibly due to lipid rearrangements (Supplementary Fig. S4 and S5). The bacteria were then incubated with various concentrations of SA, and the corresponding *D* values were determined and plotted as a function of SA concentration (Figure 3d). The resulting plot shows a clear, concentration-dependent increase in *D*, indicating enhanced lateral diffusion within the membrane with increasing SA levels. Due to solubility limitations, SA concentrations were restricted to a maximum of 0.03%, which corresponded to the concentrations showing the largest changes in growth kinetics. Since SA is a lipophilic molecule, it can readily partition into the lipid bilayer, thereby altering the organization and mobility of membrane components. This interaction is evident from changes in membrane dynamics, as indicated by a faster lipid-molecule transit time following SA treatment. To examine whether the SA-induced changes were reversible, we performed agarose pad experiments. The results showed that these effects were transient and reverted to their original state after incubation in SA-free medium (Supplementary Fig. S6).

### Viscosity measurements of the bacterial membrane using FLIM

To examine how SA affects the bacterial membrane environment, FLIM was performed using the molecular rotor BODIPY C12, which is sensitive to lipid packing and membrane viscosity and selectively partitions into the lipid membrane (Figure 4a,b). Panels in Figure 4a and b represent the intensity image, lifetime map, and the corresponding viscosity map extracted from the FLIM data for pristine cells and cells treated with 0.03% SA, respectively. Panel c shows the fluorescence lifetime decay profiles obtained from the FLIM images after applying an appropriate threshold to exclude the background signal. A noticeable decrease in fluorescence lifetime was observed after SA treatment (Figure 4c), indicating a decrease in membrane viscosity. The average viscosity (*η*) values for pristine bacteria were approximately 1200 *±* 330 cP, while for cells incubated with 0.03% SA, the *η* was around 1064 *±* 328 cP. Figure 4d represents the viscosity histograms for different SA concentrations extracted from the FLIM images using the Förster-Hoffmann equation (FH), which showed a concentration-dependent shift toward lower *η* values, consistent with the *D* trends obtained from FCS in Figure 3. To convert the *D* in Figure 3c to viscosity values, we used the Saffman–Delbrück diffusion equation (supplementary material). The *η* values obtained from the *D* map are in close agreement with those extracted from the fluorescence lifetime values in the FLIM images. Qualitatively, the results are comparable; however, the *D* map exhibits a more pronounced change after SA incubation. These results are consistent with the trends observed from the FCS measurements, although the magnitude of change detected by FCS was greater than that observed in the viscosity measurements (Fig. S8 in SM). No significant variation was observed in the lifetime of the PG layer labeled with WGA (Fig. S9 in SM), a specific PG marker, either in the presence or absence of SA, indicating that SA does not measurably alter the local environment of the PG layer.

**FIG. 4.**
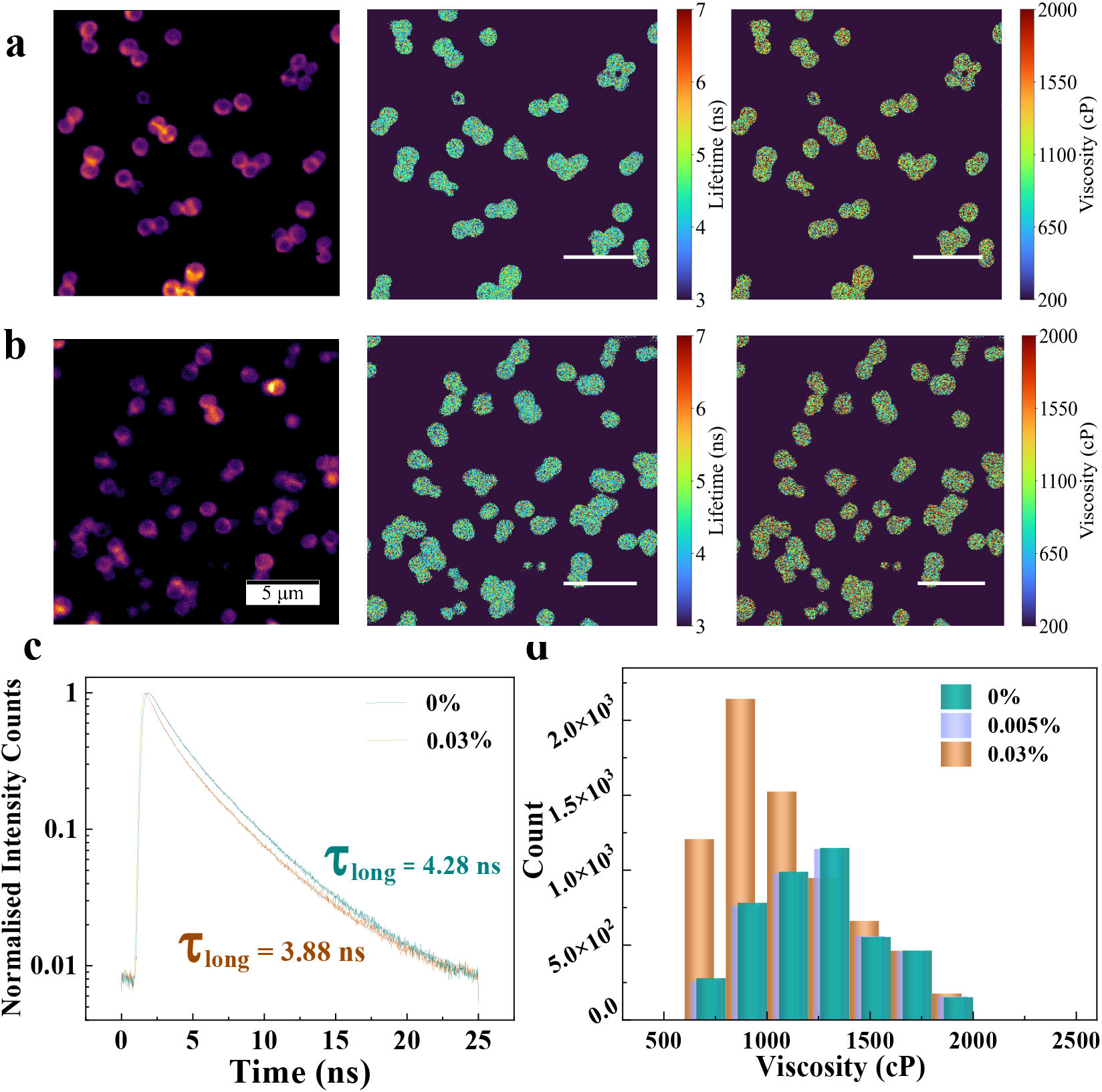
Panel (a) Intensity image, lifetime, and viscosity map for the pristine *S. epi* labelled with BODIPY C12, membrane labelling dye. (b) For the bacteria incubated with 0.03% of SA. (c) Lifetime decay profiles for pristine *S. epi* and 0.03% of SA. (d) Histograms of viscosity map for various concentrations of SA. It can be seen from the figure that viscosity histograms qualitatively follow the diffusion coefficient trend before and after incubation with SA.

### Interaction of SA with the inner membrane of *S. epi* using Martini simulations

To provide molecular insights into our experimental findings on the inner membrane (IM), we carried out CG-MD simulations with the *S. epi* membrane consisting of DAGX, PGLD and CDLX in the ratio, 45:33:22. The S. epi, membrane composition and CG parameters were obtained from a recent lipidomics and force-field parametrization study in our laboratory [47]. The lipids have headgroups of varying excluded volumes, and branched anteiso fatty acid tails with a distinct lipid tail asymmetry (C15:C20) as illustrated in Figure 5a. In contrast SA has a C18 tail with a smaller headgroup resulting in a mismatch with both lipid headgroups and hydrocarbon tails of the *S. epi* membrane. The MD simulation details are given in Appendix G. Simulations were carried out with 2, 10 and 30 SA molecules initially placed in the extracellular space. SA molecules spontaneously insert within 100 ns into the membrane. The visualization of SA trajectories with 10 and 30 SA molecules revealed that multiple SA molecules initially assembled into a micellar aggregate in the aqueous phase [10], which subsequently fused with the membrane (video in supplementary material). The density profiles for the headgroup (COOH) and terminal carbon bead of SA given in Fig. S11 of SM indicate the preferential orientation of the polar headgroup, aligning with the lipid head-groups at the lipid-water interfaces. The SA head-group density in the core of the bilayer arises from the flip-flop events of SA molecules. The center-of-mass motion of SA molecules depicted in Supplementary Fig. S13 illustrates the positioning of SA molecules in the membrane. The SA molecules sampled different regions of the bilayer interior. A fact that SA molecules undergo several flip-flop events across the leaflets is an indicative of the low free energy barrier to translocate across the core of bilayer, (Figure 5c). The potential of mean force (PMF) computed using umbrella sampling for insertion of a single SA molecule into the *S. epi* membrane is illustrated in Figure 5c. From restraint-free simulations, we calculated the mean squared displacements (MSD) for the lipids and SA to bring out the effect of SA on diffusion dynamics of lipids. Data for 2 and 30 SA molecules are given in Fig. S12 in the SM. Figure 5d illustrates the MSD for the DAGX lipid with and without SA molecules. The lateral diffusion coefficients, *D*, for all the lipids deduced from MSD fits are given in Table (supplementary material). Enhanced diffusion of membrane lipids in the presence of SA molecules (Figure 5c), is in agreement with the SA-induced increased fluidity of the IM observed in the experiments (Figure 3). We also observed an increasing trend in lipid diffusion coefficient from 5.0 *×* 10^−7^ cm^2^/s with 2 SA molecules to 5.85 *×* 10^−7^ cm^2^/s at 30 SA molecules (Table S3 in SI). This amounts to a percentage rise of 57-84% with an increasing SA concentration. This increase is similar to the enhancement observed in the experiments.

**FIG. 5.**
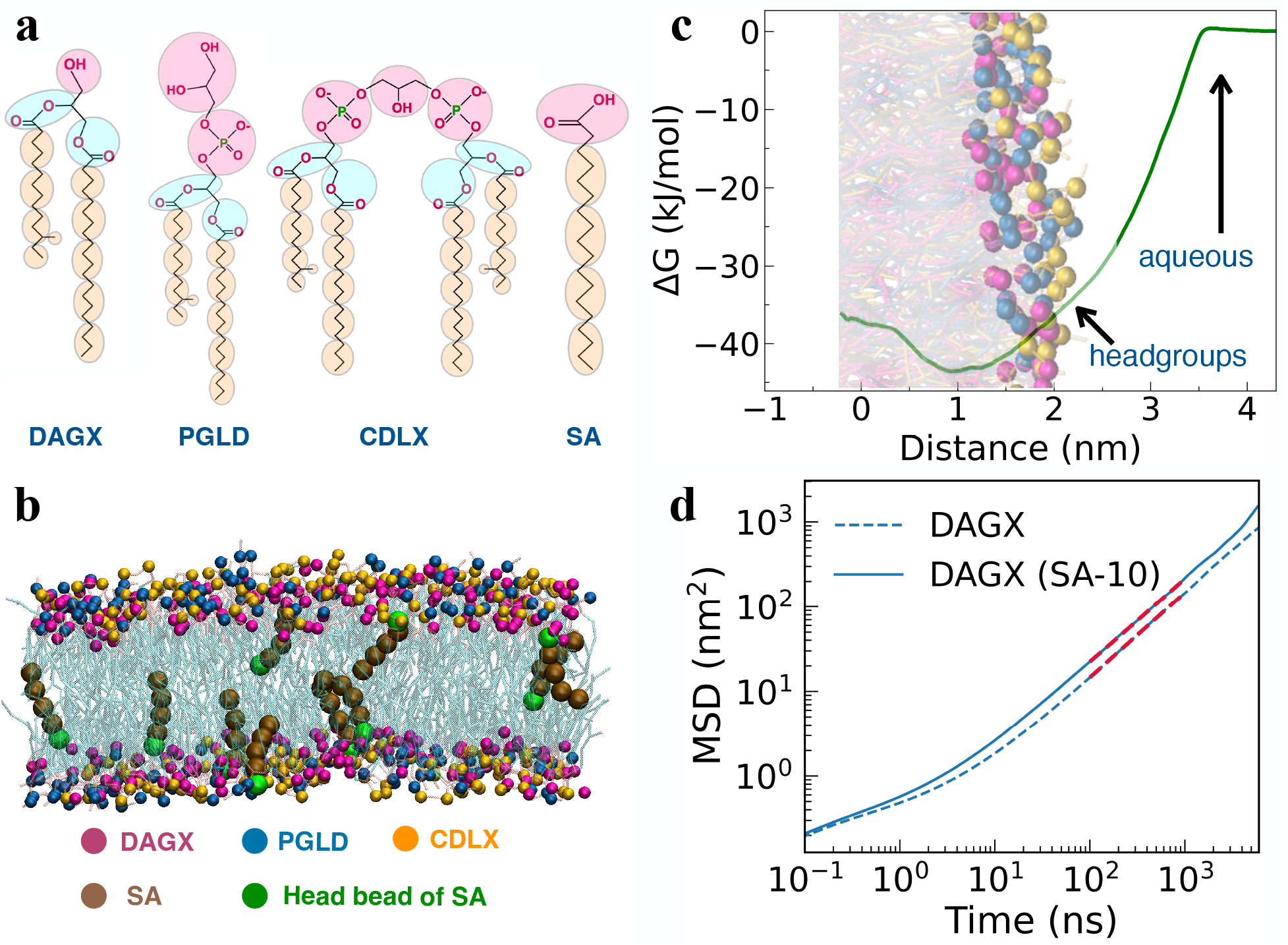
(a) Coarse-grained molecular models for *S. epi* membrane lipids, namely diacylglycerol (DAGX), phosphatidylglycerol (PGLD), and cardiolipin (CDLX) and SA molecule. (b) The membrane model of *S. epi*, with SA embedded into the membrane shown with its tail beads (brown color) and headgroup (green). (c) Profile for the potential of mean force as a function of the distance between the center-of-mass of SA and the membrane centre. (d) Mean squared displacement of DAGX lipid. Here, the solid lines correspond to the MSD for the case with 10 molecules of SA, while blue dotted lines represent the MSD for the case without SA, and red dotted lined represent the fits for MSD=4*Dt*.

### Effect of SA on the mechanical properties of the bacterial cell envelope

Exposure to antimicrobial molecules causes membrane fluidization and reduced mechanical stability in bacterial cells, and in the present study, SA treatment increases membrane dynamics while supporting bacterial growth.This raises the question of how increased lipid mobility is maintained without loss of structural integrity. To address this, we used AFM to probe the mechanical response of the outer cell envelope, which in Gram-positive bacteria is dominated by the PG layer. Contact-mode AFM was performed on *S. epi* cells under liquid conditions to examine cell morphology and to evaluate the mechanical behaviour of the bacterial envelope. Cell stiffness, one of a key mechanical properties, was quantified from force–distance (F–D) curves using the classical Hertz model [48, 49]. In the context of AFM F-D spectroscopy, according to the classic Hertz model, the indentation force (F) applied to an infinitely elastic half-space (bacterial surface) is given by

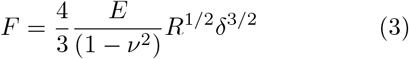

where *E* is the Young’s modulus of the bacteria, *ν* is the Poisson’s ratio of the bacterium [50], taken as 0.5 [51], *R* is the radius of the AFM tip, and *δ* is the indentation depth. From the F-D data, the Young’s modulus can be derived. Higher values correspond to a stiffer cell envelope. Since the indentation depth used for fitting (50 nm) is comparable to the reported thickness of the Gram-positive PG layer [2, 32], the measured mechanical response is dominated by the PG network more than the underlying lipid bilayer.

Figure 6a shows the height and drive amplitude images of *S. epi* bacteria in HEPES buffer. Figure 6b shows a representative F–D curve on the bacterial surface, where the major contribution comes from the cell wall, which is in the range of 30 nm thick for *S. epi* bacteria [32, 52]. The F–D curve exhibits a steeper slope for bacteria incubated with SA, indicating that indentation is more difficult compared to pristine bacteria, where the slope change is less pronounced. Young’s modulus values were extracted using the contact Hertz model described above, and the resulting histogram is shown in Figure 6c. The Young’s modulus of pristine *S. epi* cells was 1.52 MPa, which increased to 2.30 MPa (34.7%) after incubation with SA, indicating a significant enhancement in cell stiffness. The reported values for pristine *S. epi* cells are comparable to the values reported by Han et.al [32]. The Young’s modulus of the substrate with different surface coatings was also measured and found to be in the range of tens of MPa, considerably higher than that of the bacterial cell surface ((Supplementary Fig. S10). This increase in Young’s modulus suggests that the bacterial cell wall became stiffer and contributed predominantly to the observed change. To verify these results, bacteria were also adhered to high-density PLL-coated substrates, and the experiments were repeated, and the results are shown in (Supplementary Fig. S10). Qualitatively, the experimental results were consistent across different substrates, with minor variations in Young’s modulus values likely arising from substrate effects.

**FIG. 6.**
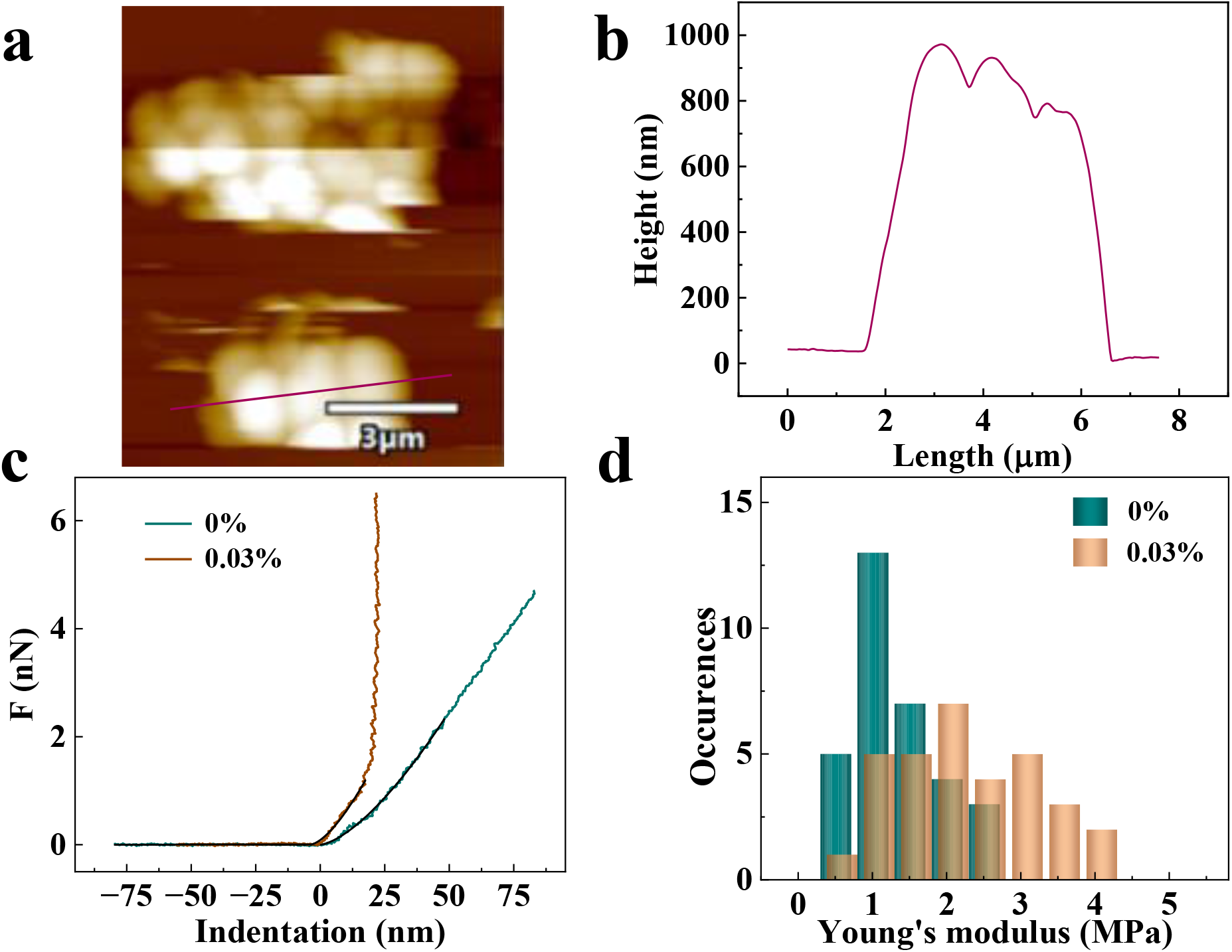
(a) AFM height image of *S. epi* bacteria. (b) Line profile corresponding to the height image extracted along the line drawn on the bacteria in (a), as indicated. (c) F–D spectroscopy curves for pristine bacteria, bacteria incubated with SA and Substrate, with fit lines shown in black. (d) Histogram of pristine *S. epi* and SA-incubated bacteria, clearly showing an increase in Young’s modulus values, indicating that SA treatment has rigidified the PG layer of the envelope compared to pristine bacteria.

**FIG. 7.**
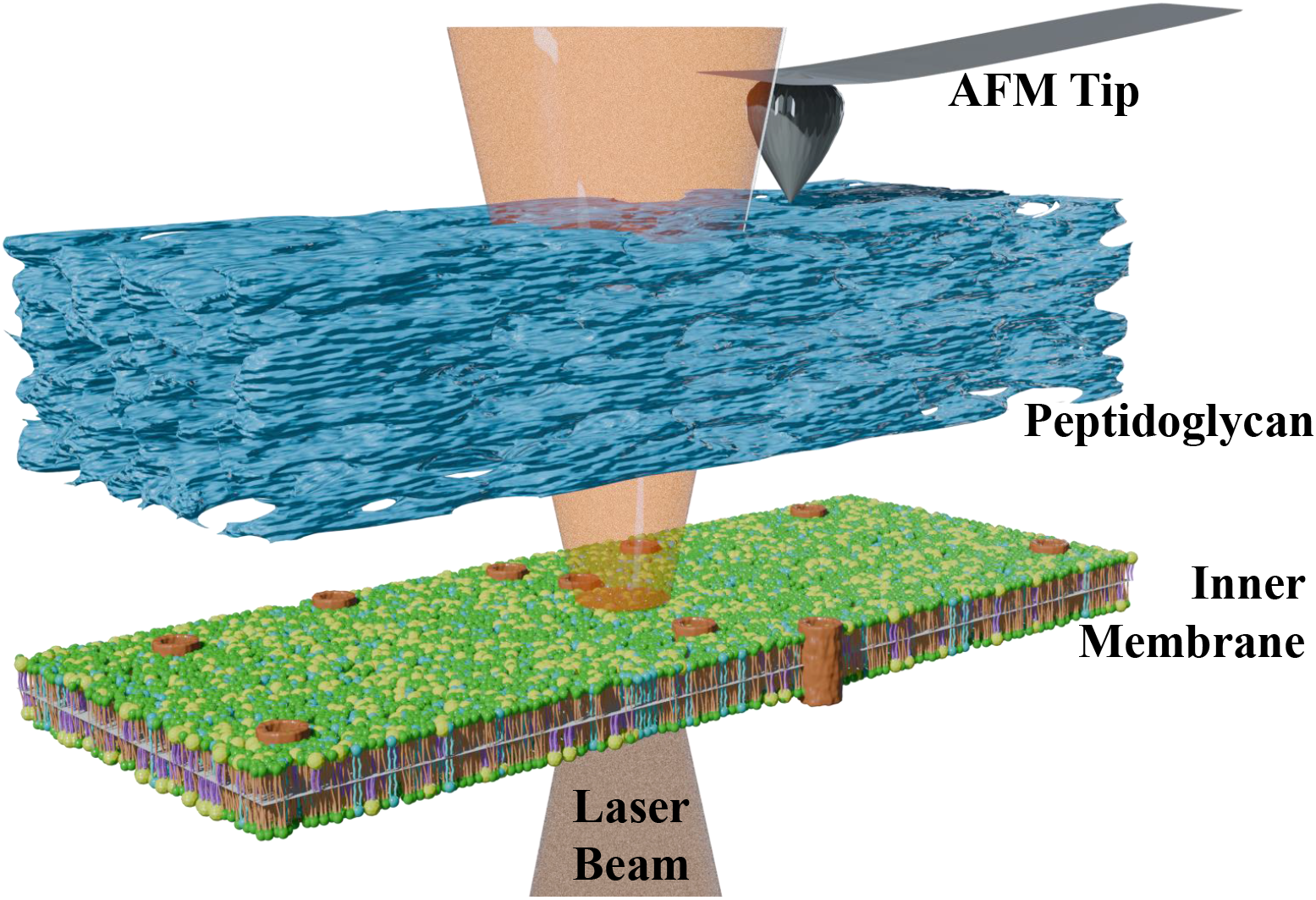
The schematic illustrates the experimental approaches used to investigate bacterial cell envelope properties. Mechanical properties of the PG layer were quantified using AFM, nano-indentation method. Lipid dynamics of the NR- and BODIPY C12–tagged IM, including the diffusion coefficient and membrane viscosity, were probed using FCS and FLIM.

## III. DISCUSSION

Growth kinetics curves of *S. epi* with and without SA were fitted using the modified Gompertz model. The analysis revealed that exposure to SA led to a significant increase in the maximum specific growth rate and an enhanced cell density increment at an early start of the log phase, indicating a potential prebiotic effect in *S. epi* [53–55]. To further understand this effect, we investigated the influence of SA on the bacterial cell envelope using a combination of dynamical and mechanical measurements. Our results show that SA incubation at sufficient levels leads to clear alterations in membrane-associated dynamics as well as changes in envelope stiffness, mainly contributed by the thick PG layer in Gram-positive bacterial membranes [2, 56], demonstrating that both structural and functional properties of the bacteria are modulated by SA. It is well understood that different classes of fatty acids interact with lipid membranes in distinct ways. Mono and poly-unsaturated fatty acids possess kinks in the acyl chains and disrupt lipid packing, which increases membrane fluidity, whereas saturated fatty acids tend to enhance membrane rigidity by promoting tighter, more ordered packing. However, these effects are concentration-dependent, as excessive incorporation of fatty acids can destabilize the bilayer and compromise membrane integrity [57, 58]. Many antimicrobials exert their bactericidal effect mainly by targeting the bacterial membrane, causing membrane disruption and fluidization [31].

In this work, we observe a similar effect when *S. epi* is incubated with SA, where membrane fluidization occurs but leads to the positive changes in growth kinetics, when molecules interact with membranes. This suggests that the IM fluidization may be part of a bacterial rearrangement mechanism in both scenarios. We have earlier shown that single surfactant molecules with varying chain lengths and degrees of saturation can translocate across the PG [10], and micellar aggregates larger than the PG void size (2-20 nm) are unable to cross the PG. Hence, SA is expected to cross the PG network of *S. epi* in their monomeric form and increase the fluidity of the underlying lipid bilayer, supporting our observations. The traditional view is that SA would reduce the fluidity of ceramides and phospholipid bilayers [59, 60]. However we observed the opposite effect, where the fluidity of the bilayer is increased in the presence of SA. Reduced mobility is associated with improved packing and this occurs when SA is hydrophobically matched with the underlying membrane lipids. In the case of *S*.*epi*, both the SA hydrocarbon tails and the headgroup excluded volume are mismatched (Figure 5a), as a result SA is laterally mobile, freely sampling the lateral extent of the bilayer in the x-y plane. Additionally, SA undergoes a large number of flip-flop events across the bilayer (Supplementary Fig. S13).This leads to the increased mobility in the *S. epi* IM lipids.

To evaluate whether alterations in the lipid membrane are linked to changes in the mechanical properties of the PG, we measured the Young’s modulus using AFM force–distance spectroscopy. An increase in Young’s modulus was observed after incubation with SA, suggesting enhanced stiffness of the PG layer. This stiffening may result from enhanced peptide cross-linking, a denser PG network with smaller effective pores, increased glycan strand density, or changes in the mechanical interactions between the membrane and the PG layer, all of which are known to modulate the elastic modulus of bacterial cell walls [36, 52, 61]. Furthermore, AFM measurements by Bailey et al. showed that the stiffness of isolated cell wall of *sacculi* is comparable to that of intact cells, indicating that cell stiffness measured by AFM is dominated by the mechanics of the PG layer in Gram-positive bacteria [36].

Our results, therefore, suggest that SA may influence the cell envelope through a distinct mechanism compared to conventional antibiotics. Although the PG exhibits enhanced rigidity upon SA treatment, the lipid membrane was also fluidized, as evident from the increased lateral diffusion and decreased viscosity data. Overall, the growth kinetics curves suggest a possible prebiotic effect of SA, which was further analyzed in detail using the Gompertz model. Lipid dynamics measurements clearly show membrane fluidization upon SA treatment, in agreement with the MD simulation results. These altered lipid dynamics may trigger subsequent biochemical responses [40] that lead to stiffening of the PG layer, which in turn modifies the bacterial cell envelope properties and contributes to the observed prebiotic effects.

We conclude the discussion by drawing a connection between the observed increase in membrane fluidity and the enhanced PG rigidity in the presence of SA in Gram-positive bacteria. There are several reports that connect increased membrane fluidity and lateral membrane organization to assist PG synthesis [39, 62]. In an early study [63] addition of n-butanol and other higher alkanes to *S. aureus* was found to upregulate enzyme activity in the PG synthesis pathway – attributed primarily to the increased membrane fluidity. A more recent and compelling view that connects the fluidity of the lipid membrane environment is related to lipid II, a key player in the cell wall synthesis pathway [64].

Lipid II is made up of a long hydrophobic tail attached to a disaccharide-pentapetide group made up of N-acetlyglucasamine and N-acetylmuramic acid with a pentapeptide stem, the primary building block for the cell wall. Lipid II is enzymatically synthesized in the cytosolic side of the lipid bilayer and transported across to the upper leaflet, where the penicillin binding protein (PBP) catalyzes glycan strand polymerization and the cross-linking between glycan chains with peptide linkages to complete the synthesis [42, 65]. It has been hypothesized that PG synthesis can potentially be upregulated in a more fluid-like IM environment that assists transport of lipid II and other downstream enzymatic pathways in the IM [64]. The upregulation of PG synthesis can result in increased stiffness, as observed in this study. Given the above connects with membrane fluidity and PG synthesis pathways, our study provides further and direct support to the view that IM fluidity and cell wall synthesis are intrinsically connected, with increased fluidity correlating positively with enhanced cell wall stiffness. Finally, enhanced membrane fluidity may also facilitate the transport of nutrients required for cell wall synthesis.

## IV. APPENDIX

*S. epi*, ATCC 12228 was obtained from the American Type Culture Collection and cultured overnight in tryptic soy broth (TSB; Difco™) at 37^*◦*^C with continuous shaking. The bacterial suspension was adjusted to an optical density at 620 nm (OD_620_) using a spectrophotometer to achieve a final concentration of approximately 10^8^ CFU/mL. SA was purchased from Sigma-Aldrich, and a stock solution of 10 % (w/v) was prepared in a mixture of DMSO, Tween 80 and deionized water in the growth (nutrient) media in a ratio of 80:10:10. Working solutions were freshly prepared by diluting the stock as required for the experiments. Poly-L-lysine (PLL) and Vectabond, purchased from Sigma Aldrich and Vector Laboratories, respectively, were used to immobilize the cells during imaging. Nile red, a fluorescent marker, was used for labelling and was procured from Sigma Aldrich as well. Nile red was employed for imaging and FCS.

### A. Bacterial cultures

To initiate the primary culture of the cells, a sterile loop or inoculation needle was used to pick up a portion of cells from a single colony TSA plate, which was then inoculated into 5 mL of TSB medium. The culture was incubated overnight at 37°C with continuous shaking with an rpm of 180 in the incubator. For concentration-dependent microscopy studies, secondary cultures were grown in the presence of SA to an OD of 0.5 before being used for imaging and other measurements.

### B. Kinetic measurements

Working solutions of SA were freshly prepared by diluting the stock solution to the required concentrations. For growth studies, a 100-fold dilution of the primary inoculum grown overnight was added to fresh TSB medium prepared at a 1:10 dilution in DI water. The bacterial suspension in each well was adjusted to a final concentration of approximately 10^5^ cfu/ml. Absorbance was measured at OD_620_ using a spectrophotometer (Tecan, Infinite 200 PRO) as an indicator of bacterial cell density. The cultures were incubated at 37°C in 96-well plates with intermittent shaking (every 15 minutes at 180 rpm), and absorbance data were collected continuously over 24 hours at 15-minute intervals. Due to solubility limitations of SA in the growth medium, experiments were performed up to a maximum SA concentration of 0.03%. OD_620_ values were plotted as a function of time for pristine samples and for various concentrations of SA, together with positive and negative controls.

### C. Fluorescence correlation spectroscopic (FCS) measurements

*S. epi* cells were grown to late log phase at 37^*◦*^C, labelled with 0.5 to 1 *µ*g/ml of NR and washed multiple times with PBS to remove excess dye. The bacteria were spread on a PLL coated coverlip to promote adhesion to the substrate, followed by additional PBS washes to remove non-adherent cells. Imaging was performed using an SP5 Leica confocal laser scanning microscope (Leica Microsystems, Germany) integrated with a STEDYCON super-resolution module (Abberior Instruments GmbH). NR labels the bacterial membrane, and FCS was performed in the polar region of the cells. The resulting histograms are compilations of approximately 60–80 readings taken from 8–10 bacteria. All experiments were conducted in triplicate, and the data are presented as the mean *±* standard deviation. NR was excited using a 561 laser, and the emission spectrum was recorded with a 575–625 nm avalanche photodiode detector. Intensity data were correlated using PicoQuant SymPhoTime software. The intensity traces were converted into autocorrelation profiles using the following calculations. In FCS, the autocorrelation function G(*τ*) from the intensity signal I(t) measured under the microscope is calculated using,

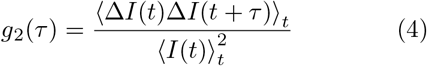

where *<>*_*t*_, denotes that the physical quantity was averaged over the time variable t. The correlation data were normalized and fitted to a single or multi-component 2D diffusion equation, the following equation:

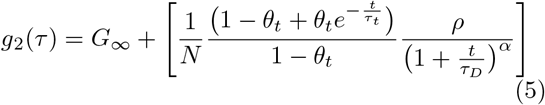

where *G*_∞_ is the background of the curve where the autocorrelation decays to 0. *α* is the anomaly parameter, which is 1 for Brownian diffusion and deviates from 1 in the case of the sub-diffusive or super-diffusive nature of diffusion. *θ*_*t*_ is the fraction of molecules that are present in the triplet state, *τ*_*t*_ is the relaxation time in the triplet state, *ρ* is the fraction of diffusing molecules, which is unity in these fits, and *τ*_*D*_ is the diffusion time. The Quickfit software was used to extract the *τ*_*D*_ from Equation 5. The anomaly parameter *α* values were approximately 0.9 *±* 0.1 for our profiles, suggesting Brownian diffusion. Once the *τ*_*D*_ values are known, the diffusivity *D* can be evaluated using the following equation

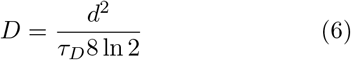

where *d* represents the full width at half maximum (FWHM) of the Gaussian beam. *τ*_*D*_ is the diffusion time, and D is the diffusion coefficient of the NR molecules in the lipid environment. The model autocorrelation profiles with fits are given in Figure 3.

### D. Atomic force microscopy (AFM)

13 mm glass coverslips were cleaned using a 1–2% SDS solution in a 37 kHz operating sonicator for 10 minutes, thoroughly rinsed with Milli-Q water, followed by rinsing with ethanol, dried with Nitrogen and subsequently plasma cleaned for 2 minutes at 70% power [66]. The cleaned coverslips were then coated with Vectabond:Acetone (50:1) for 5 minutes and thoroughly rinsed with water. After drying with Nitrogen, bacterial samples were spread onto the Vectabond-coated coverslips and immobilized for experimental use. Imaging was performed in the contact mode under liquid conditions with the sample immersed in HEPES buffer, with the Asylum AFM incorporated with an Olympus inverted optical microscope. Images were taken with the tip that has a tip frequency of 17 KHz, with the spring constant of 0.08 N/m and the set point of 40 nN. Calibration of the cantilever was done using a DPPC bilayer of known Young’s modulus. The effective Young’s modulus was calculated using the Hertz–Sneddon model with a paraboloid tip shape, and a Poisson ratio of 0.5 was used for the bacterial samples.

### E. Fluorescence life time imaging (FLIM)

0.5 OD bacterial cultures, with and without SA, were labelled with BODIPY C12, a membrane tagging dye, for 2 hours at a final concentration of 200 nM [67], as this concentration does not affect bacterial growth. Upon labelling, cells were pelleted and resuspended in PBS buffer (pH 7.0). The bacterial suspension was spread on a slide coated with 0.1% PLL to promote adhesion, then imaged usingFLIM with a time-correlated single-photon counting unit (TCSPC) (SPC-830; Becker & Hickl, Berlin, Germany) integrated into a confocal microscope (TCS SP5; Leica Microsystems, Wetzlar, Germany). Excitation was performed at 488 nm, and emission was collected in the 500–550 nm range corresponding to BODIPY C12 fluorescence. The dye concentration chosen (200 nM) provided an optimal signal-to-noise ratio for analysis. The images were processed using Python to convert lifetime maps into viscosity maps using the Förster-Hoffmann equation. The regions of interest (ROIs) were selected by applying a threshold to remove background noise, and the viscosity values were extracted for the entire image frame to generate histograms. For selected ROIs in the FLIM images, the corresponding TCSPC lifetime decays were extracted and fitted using a two-component decay model, from which the *τ*_*avg*_ was determined. The membrane viscosity was then calculated using the formula given below.

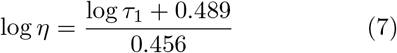

In the above equation, *η* denotes the viscosity in cP, and *τ*_1_ denotes the longer-life component. The constants 0.489 and 0.456 were calculated from the calibration curve using a previously reported viscosity-lifetime calibration equation, obtained by measuring the fluorescence decays of BODIPY C10 in methanol/glycerol mixtures of known viscosity (*η*) [68] (Supplementary Fig. S7).

To investigate the lifetime–viscosity trends in greater detail, fluorescence decay curves were extracted from the selected ROI and analyzed using a biexponential fitting model. The quality of the fits was evaluated using the chi-squared parameter (*χ*^2^), which was approximately 0.99 for the fits shown in Figure 4c, indicating a good fit. ROIs were chosen through thresholding to ensure that only regions with a good signal-to-noise ratio were included in the analysis. The fluorescence decay profiles were then fitted to a biexponential model, which is mathematically expressed.

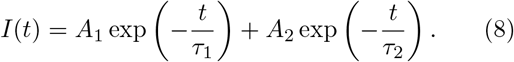

Here, *A*_1_ and *A*_2_ represent the amplitudes of the two exponential components, while *τ*_1_ and *τ*_2_ denote their respective lifetimes. Once these parameters are determined, the average lifetime (*τ*_avg_) can be calculated using the weighted average method, as shown in Equation 9.

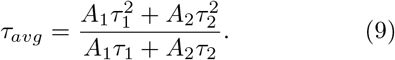

We have used a longer component of the lifetime to calculate the viscosity.

### G. MD simulations

We have carried out MD simulations on *S. epi* membrane composed of diacylglycerol (45 mol %), phosphatidylglycerol (33 mol %) and cardiolipin (22 mol %). The lipidomics study has been carried out to determine the lipid compositions of the S. epi ATCC 12228 bacterial strain using a proprietary Shotgun Lipidomics platform, and its details are given in a separate study on Gram-positive bacteria [47]. The CG models for these lipids (Figure 5) were developed using the Martini-3 force field, and the CG membrane model was validated by comparing structural properties of the membrane with atomistic simulations using the CHARMM36 force field. The membrane was composed of 232 DAGX, 168 PGLD and 112 CDLX lipids. The membrane charges were neutralized by adding potassium ions. The equations of motion were integrated using a leapfrog algorithm with a time step of 20 fs. The non-bonded van der Waals interactions (Lennard Jones potential) were computed within a cutoff radius of 1.1 nm. The long range coulomb interactions were evaluated using the reaction field method with a relative dielectric constant of 15 and a Coulomb cutoff of 1.1 nm. The system temperature was maintained using v-rescale thermostat [69] coupled at every 1 ps. The Parrinello Rahman barostate [70] was used to control the pressure in semi-isotropic mode with coupling constant 12 ps and compressibility 3 *×* 10^−4^ bar^−1^. The umbrella sampling simulations were performed to compute the interaction of an SA molecule with the *S. epi* membrane. The membrane system was equilibrated for 10 microseconds at 310 K and 1 bar, after which the potential of mean force (PMF) was calculated. A total of 45 umbrella windows with a spacing of 0.1 nm were used, extending over a distance of ∼ 4.5 nm from the bulk water phase to the centre of the membrane. The initial configurations for the umbrella sampling simulations were generated from a steered MD simulation, and each umbrella window was subsequently simulated for 200 ns in the production run to obtain a smooth and converged free energy profile. In addition, 10 microsecond long restraint-free simulations were carried out with 2, 10, and 30 SA molecules to demonstrate the spontaneous insertion of SA into the lipid membrane and its modulation in lipid diffusion at low, medium and high concentrations of SA.

## ACKNOWLEDGMENTS

We thank Unilever Research and Development (Bangalore, India) for funding this research (Project code: 99537). The authors thank IISc for support under the Ministry of Education (MoE) and Institute of Excellence (IoE) grants. Some authors acknowledge support from the Department of Science and Technology (DST) through the Funding for Improvement of Science and Technology (FIST) grant. KGA thanks Arul Mozhy Varman for several insightful discussions. SJJ acknowledges financial support from the University Grants Commission (UGC) through a UGC Research Fellowship.

## References

[1] G. Rani and I. Patri, Modeling heterogeneities in the crosslinked bacterial sacculus, Physical Review Research 2, 013090 (2020).

[2] T. J. Silhavy, D. Kahne, and S. Walker, The bacterial cell envelope, Cold Spring Harbor perspectives in biology 2, a000414 (2010).

[3] M. Rajagopal and S. Walker, Envelope structures of gram-positive bacteria, Protein and sugar export and assembly in Gram-positive bacteria, 1 (2016).

[4] H. Hauser, M. Phillips, and M. Stubbs, Ion permeability of phospholipid bilayers, Nature 239, 342 (1972).

[5] D. W. Deamer and J. Bramhall, Permeability of lipid bilayers to water and ionic solutes, Chemistry and physics of lipids 40, 167 (1986).

[6] J. Sun, S. T. Rutherford, T. J. Silhavy, and K. C. Huang, Physical properties of the bacterial outer membrane, Nature Reviews Microbiology 20, 236 (2022).

[7] W. N. Konings, S.-V. Albers, S. Koning, and A. J. Driessen, The cell membrane plays a crucial role in survival of bacteria and archaea in extreme environments, Antonie Van Leeuwenhoek 81, 61 (2002).

[8] N. Marín-Medina, D. A. Ramírez, S. Trier, and C. Leidy, Mechanical properties that influence antimicrobial peptide activity in lipid membranes, Applied microbiology and biotechnology 100, 10251 (2016).

[9] G. K. Auer and D. B. Weibel, Bacterial cell mechanics, Biochemistry 56, 3710 (2017).

[10] P. Sharma, R. Vaiwala, S. Parthasarathi, N. Patil, A. Verma, M. Waskar, J. S. Raut, J. K. Basu, and K. G. Ayappa, Interactions of surfactants with the bacterial cell wall and inner membrane: revealing the link between aggregation and antimicrobial activity, Langmuir 38, 15714 (2022).

[11] B. Mostofian, T. Zhuang, X. Cheng, and J. D. Nickels, Branched-chain fatty acid content modulates structure, fluidity, and phase in model microbial cell membranes, The Journal of Physical Chemistry B 123, 5814 (2019).

[12] Y.-Y. Chang and J. E. Cronan, Membrane cyclopropane fatty acid content is a major factor in acid resistance of escherichia coli, Molecular microbiology 33, 249 (1999).

[13] J. J. Baldassare, K. B. Rhinehart, and D. F. Silbert, Modification of membrane lipid: physical properties in relation to fatty acid structure, Biochemistry 15, 2986 (1976).

[14] A. P. Desbois and V. J. Smith, Antibacterial free fatty acids: activities, mechanisms of action and biotechnological potential, Applied microbiology and biotechnology 85, 1629 (2010).

[15] J. J. Kabara, D. M. Swieczkowski, A. J. Conley, and J. P. Truant, Fatty acids and derivatives as antimicrobial agents, Antimicrobial agents and chemotherapy 2, 23 (1972).

[16] E. Obukhova and S. Murzina, Mechanisms of the antimicrobial action of fatty acids: A review, Applied Biochemistry and Microbiology 60, 1035 (2024).

[17] J. J. Kabara, Antimicrobial agents derived from fatty acids, Journal of the American Oil Chemists’ Society 61, 397 (1984).

[18] K. Fritsche, Fatty acids as modulators of the immune response, Annu. Rev. Nutr. 26, 45 (2006).

[19] A. P. S. Caldas, D. M. U. Rocha, J. Bressan, and H. H. M. Hermsdorff, Dietary fatty acids as nutritional modulators of sirtuins: a systematic review, Nutrition reviews 79, 235 (2021).

[20] Y. Endo, S. Kamisada, K. Fujimoto, and T. Saito, Trans fatty acids promote the growth of some lactobacillus strains, The Journal of General and Applied Microbiology 52, 29 (2006).

[21] P.E. Kankaanpää, S. J. Salminen, E. Isolauri, and Y. K. Lee, The influence of polyunsaturated fatty acids on probiotic growth and adhesion, FEMS microbiology letters 194, 149 (2001).

[22] M. K. Mitchell and M. Ellermann, Long chain fatty acids and virulence repression in intestinal bacterial pathogens, Frontiers in Cellular and Infection Microbiology 12, 928503 (2022).

[23] K. T. Yuyama, M. Rohde, G. Molinari, M. Stadler, and W.-R. Abraham, Unsaturated fatty acids control biofilm formation of staphylococcus aureus and other gram-positive bacteria, Antibiotics 9, 788 (2020).

[24] J.-S. Choi, N.-H. Park, S.-Y. Hwang, J. H. Sohn, I. Kwak, K. K. Cho, and I. S. Choi, The antibacterial activity of various saturated and unsaturated fatty acids against several oral pathogens, Journal of Environmental Biology 34, 673 (2013).

[25] Y.-M. Zhang and C. O. Rock, Membrane lipid homeostasis in bacteria, Nature Reviews Microbiology 6, 222 (2008).

[26] A. H. Cheung Lam, N. Sandoval, R. Wadhwa, J. Gilkes, T. Q. Do, W. Ernst, S.-M. Chiang, S. Kosina, H. Howard Xu, G. Fujii, et al., Assessment of free fatty acids and cholesteryl esters delivered in liposomes as novel class of antibiotic, BMC Research Notes 9, 337 (2016).

[27] E. P. Ivanova, S. H. Nguyen, Y. Guo, V. A. Baulin, H. K. Webb, V. K. Truong, J. V. Wandiyanto, C. J. Garvey, P. J. Mahon, D. E. Mainwaring, et al., Bactericidal activity of self-assembled palmitic and stearic fatty acid crystals on highly ordered pyrolytic graphite, Acta Biomaterialia 59, 148 (2017).

[28] M. Schoeler, S. Ellero-Simatos, T. Birkner, J. Mayneris-Perxachs, L. Olsson, H. Brolin, U. Loeber, J. D. Kraft, A. Polizzi, M. Martí-Navas, et al., The interplay between dietary fatty acids and gut microbiota influences host metabolism and hepatic steatosis, Nature Communications 14, 5329 (2023).

[29] S. Parthasarathi, A. Chaudhury, A. Swain, and J. K. Basu, Microscopy insights as an invaluable tool for studying antimicrobial interactions with bacterial membranes, ChemistrySelect 10, e02345 (2025).

[30] P. Sharma, S. Parthasarathi, N. Patil, M. Waskar, J. S. Raut, M. Puranik, K. G. Ayappa, and J. K. Basu, Assessing barriers for antimicrobial penetration in complex asymmetric bacterial membranes: A case study with thymol, Langmuir 36, 8800 (2020).

[31] S. Parthasarathi, A. Chaudhury, J. K. Basu, R. Yadav, and D. K. Saini, Unraveling direct correlations between membrane nanodomain reorganization and antimicrobial resistance evolution in bacterial cells, PRX Life 3, 023017 (2025).

[32] R. Han, W. Vollmer, J. D. Perry, P. Stoodley, and J. Chen, Simultaneous determination of the mechanical properties and turgor of a single bacterial cell using atomic force microscopy, Nanoscale 14, 12060 (2022).

[33] R. Han, X.-Q. Feng, W. Vollmer, P. Stoodley, and J. Chen, Deciphering the adaption of bacterial cell wall mechanical integrity and turgor to different chemical or mechanical environments, Journal of Colloid and Interface Science 640, 510 (2023).

[34] M. Mathelié-Guinlet, A. T. Asmar, J.-F. Collet, and Y. F. Dufrêne, Lipoprotein lpp regulates the mechanical properties of the e. coli cell envelope, Nature communications 11, 1789 (2020).

[35] E. R. Rojas and K. C. Huang, Regulation of microbial growth by turgor pressure, Current Opinion in Microbiology 42, 62 (2018).

[36] R. G. Bailey, R. D. Turner, N. Mullin, N. Clarke, S. J. Foster, and J. K. Hobbs, The interplay between cell wall mechanical properties and the cell cycle in staphylococcus aureus, Biophysical journal 107, 2538 (2014).

[37] C. C. Perry, M. Weatherly, T. Beale, and A. Randriamahefa, Atomic force microscopy study of the antimicrobial activity of aqueous garlic versus ampicillin against escherichia coli and staphylococcus aureus, Journal of the Science of Food and Agriculture 89, 958 (2009).

[38] G. Francius, O. Domenech, M. P. Mingeot-Leclercq, and Y. F. Dufrêne, Direct observation of staphylococcus aureus cell wall digestion by lysostaphin, Journal of bacteriology 190, 7904 (2008).

[39] K. Kurita, F. Kato, and D. Shiomi, Alteration of membrane fluidity or phospholipid composition perturbs rotation of mreb complexes in escherichia coli, Frontiers in molecular biosciences 7, 582660 (2020).

[40] A. Balasubramaniam, P. Adi, D. T. Tra My, S. Keshari, R. Sankar, C.-L. Chen, and C.-M. Huang, Repurposing inci-registered compounds as skin prebiotics for probiotic staphylococcus epidermidis against uv-b, Scientific reports 10, 21585 (2020).

[41] P. Loskill, P. M. Pereira, P. Jung, M. Bischoff, M. Herrmann, M. G. Pinho, and K. Jacobs, Reduction of the peptidoglycan crosslinking causes a decrease in stiffness of the staphylococcus aureus cell envelope, Biophysical journal 107, 1082 (2014).

[42] E. Sauvage, F. Kerff, M. Terrak, J. A. Ayala, and P. Charlier, The penicillin-binding proteins: structure and role in peptidoglycan biosynthesis, FEMS microbiology reviews 32, 234 (2008).

[43] Y. M. G. Montes, E. R. V. Calle, S. G. S. Terán, M. R. C. García, J. C. R. Nájera, and M. R. L. Vera, Growth kinetics of lactococcus lactis and lactobacillus casei in liquid culture medium containing as prebiotics inulin or fructose, Journal of the Science of Food and Agriculture 104, 1258 (2024).

[44] J. Wang and X. Guo, The gompertz model and its applications in microbial growth and bioproduction kinetics: Past, present and future, Biotechnology Advances 72, 108335 (2024).

[45] M. H. Zwietering, I. Jongenburger, F. M. Rombouts, and K. van ‘t Riet, Modeling of the bacterial growth curve, Applied and Environmental Microbiology 56, 1875 (1990), https://journals.asm.org/doi/pdf/10.1128/aem.56.6.1875-1881.1990.

[46] F. Nazzaro, F. Coppola, F. Fratianni, and R. Coppola, Fatty acids as prebiotics and their role in antibiofilm activity, Antibiotics 15, 57 (2026).

[47] R. Vaiwala, M. Waskar, and K. G. Ayappa, Model membranes for Gram-positive bacterial strains: Assessing structural and small molecule interaction differences, In Preparation (2026).

[48] E. K. Dimitriadis, F. Horkay, J. Maresca, B. Kachar, and R. S. Chadwick, Determination of elastic moduli of thin layers of soft material using the atomic force microscope, Biophysical journal 82, 2798 (2002).

[49] N. I. Abu-Lail and T. A. Camesano, The effect of solvent polarity on the molecular surface properties and adhesion of escherichia coli, Colloids and Surfaces B: Biointerfaces 51, 62 (2006).

[50] A. Touhami, B. Nysten, and Y. F. Dufrêne, Nanoscale mapping of the elasticity of microbial cells by atomic force microscopy, Langmuir 19, 4539 (2003).

[51] V. Lulevich, Y.-P. Shih, S. H. Lo, and G.-y. Liu, Cell tracing dyes significantly change single cell mechanics, The Journal of Physical Chemistry B 113, 6511 (2009), pMID: 19366241, 10.1021/jp8103358.

[52] W. Vollmer, D. Blanot, and M. A. De Pedro, Peptidoglycan structure and architecture, FEMS microbiology reviews 32, 149 (2008).

[53] E. Gomaa, Effect of prebiotic substances on growth, fatty acid profile and probiotic characteristics of lactobacillus brevis nm101-1, Microbiology 86, 618 (2017).

[54] S. You, Y. Ma, B. Yan, W. Pei, Q. Wu, C. Ding, and C. Huang, The promotion mechanism of prebiotics for probiotics: A review, Frontiers in Nutrition 9, 1000517 (2022).

[55] J. A. Van Loo, Prebiotics promote good health: the basis, the potential, and the emerging evidence, Journal of clinical gastroenterology 38, S70 (2004).

[56] H. H. Tuson, G. K. Auer, L. D. Renner, M. Hasebe, C. Tropini, M. Salick, W. C. Crone, A. Gopinathan, K. C. Huang, and D. B. Weibel, Measuring the stiffness of bacterial cells from growth rates in hydrogels of tunable elasticity, Molecular microbiology 84, 874 (2012).

[57] X. Yang, W. Sheng, G. Y. Sun, and J. C.-M. Lee, Effects of fatty acid unsaturation numbers on membrane fluidity and α-secretase-dependent amyloid precursor protein processing, Neurochemistry international 58, 321 (2011).

[58] A. I. Tyler, J. L. Greenfield, J. M. Seddon, N. J. Brooks, and S. Purushothaman, Coupling phase behavior of fatty acid containing membranes to membrane bio-mechanics, Frontiers in Cell and Developmental Biology 7, 187 (2019).

[59] K. Hac-Wydro and P. Wydro, The influence of fatty acids on model cholesterol/phospholipid membranes, Chemistry and physics of lipids 150, 66 (2007).

[60] D. Altun, P. Larsson, C. A. Bergström, and S. Hossain, Molecular dynamics simulations of lipid composition and its impact on structural and dynamic properties of skin membrane, Chemistry and Physics of Lipids 265, 105448 (2024).

[61] L. Pasquina-Lemonche, J. Burns, R. Turner, S. Kumar, R. Tank, N. Mullin, J. Wilson, B. Chakrabarti, P. Bullough, S. Foster, and J. Hobbs, The architecture of the gram-positive bacterial cell wall, Nature 582, 294—297 (2020).

[62] N. Som and M. Reddy, Cross-talk between phospholipid synthesis and peptidoglycan expansion by a cell wall hydrolase, Proceedings of the National Academy of Sciences 120, e2300784120 (2023).

[63] P. P. Lee, W. A. Weppner, and F. C. Neuhaus, Initial membrane reaction in peptidoglycan synthesis. perturbation of lipid-phospho-nacetylmuramyl-pentapeptide translocase interactions by n-butanol, Biochimica et Biophysica Acta (BBA)-Biomembranes 597, 603 (1980).

[64] A. García-Heredia, Plasma membrane-cell wall feedback in bacteria, Journal of Bacteriology 205, e00433 (2023).

[65] S. Garde, H. Selvaraj, A. Chandramouli, G. S. Reddy, D. Bahety, P. K. Chodisetti, S. S. Kamat, and M. Reddy, A conserved editing mechanism for the fidelity of bacterial cell wall biosynthesis, Proceedings of the National Academy of Sciences 122, e2505676122 (2025).

[66] G. Benn, I. V. Mikheyeva, P. G. Inns, J. C. Forster, N. Ojkic, C. Bortolini, M. G. Ryadnov, C. Kleanthous, T. J. Silhavy, and B. W. Hoogenboom, Phase separation in the outer membrane of escherichia coli, Proceedings of the National Academy of Sciences 118, e2112237118 (2021).

[67] J. T. Mika, A. J. Thompson, M. R. Dent, N. J. Brooks, J. Michiels, J. Hofkens, and M. K. Kuimova, Measuring the viscosity of the escherichia coli plasma membrane using molecular rotors, Biophysical journal 111, 1528 (2016).

[68] P.-H. Chung, Advanced fluorescence lifetime imaging and spectroscopy techniques for biological samples, Ph.D. thesis, King’s College London (University of London) (2012).

[69] G. Bussi, D. Donadio, and M. Parrinello, Canonical sampling through velocity rescaling, J. Chem. Phys. 126, 014101 (2007).

[70] M. Parrinello and A. Rahman, Polymorphic transitions in single crystals: A new molecular dynamics method, J. Appl. Phys. 52, 7182 (1981).

