## Supplementary material for "Stearic acid enhances membrane fluidization and peptidoglycan stiffness to promote stability of Gram-positive bacteria"

### Stearic acid enhances cell envelope mechanics and promotes stability in Gram-positive bacteria

### Growth kinetic studies

For the growth kinetics study, a 1:10 dilution of Tryptic Soy Broth (TSB, Sigma Aldrich) was prepared. A 5% stock solution of stearic acid (SA) from Sigma Aldrich (Cat.log.number S4751) was prepared in a solvent mixture of DMSO:Tween 80:Water. Working concentrations of SA at 0.005%, 0.01%, 0.02%, 0.025 % and 0.03% were then prepared in 10 mL of the diluted TSB medium. An overnight-grown bacterial culture of *Staphylococcus epidermidis* (*S. epi*)was used as the inoculum to initiate the growth kinetics experiments. Growth curves were monitored over a 24-hour period, with optical density (OD) measurements recorded at 15-minute intervals. As shown in Figure S1a, an increase in SA concentration led to a corresponding increase in OD, along with a prolonged stationary phase. Due to the solubility limitations of SA in the growth medium, experiments could be carried out reliably up to a concentration of 0.03

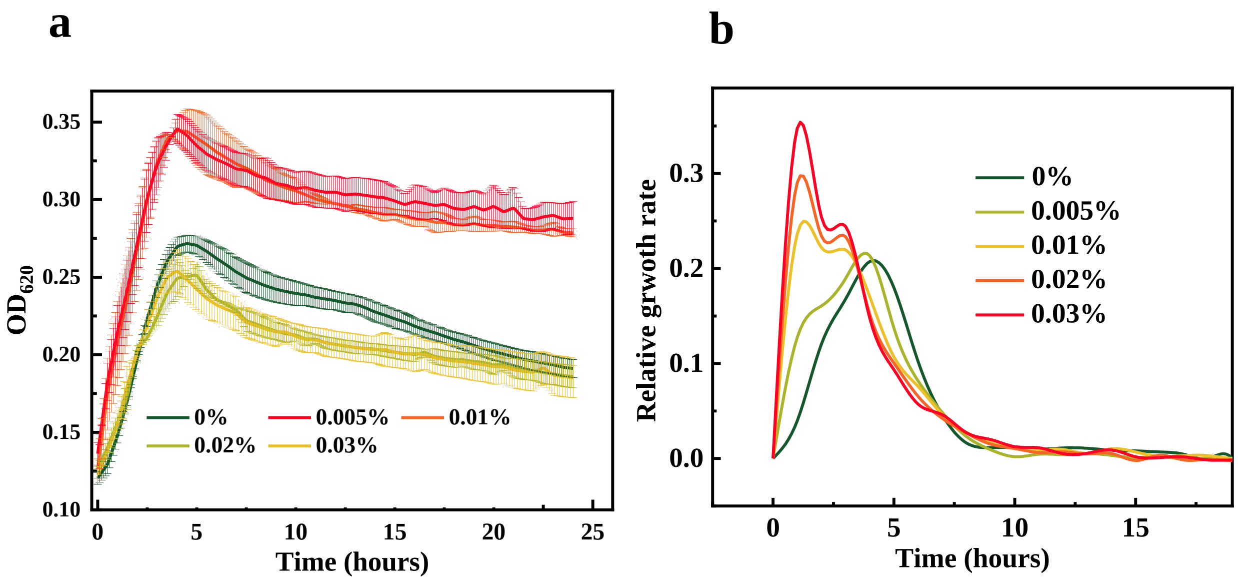


**Figure S1:** **(a)** Growth curves showing optical density ($OD_{620}$) versus time for bacterial cultures incubated with different concentrations of SA. **(b)** Relative change in growth rate over time. These data from an independent experimental set confirm the trends of enhanced growth and a prolonged stationary phase presented in the main manuscript.

To quantify the bacterial growth rate, the slopes of the late log phase were extracted for

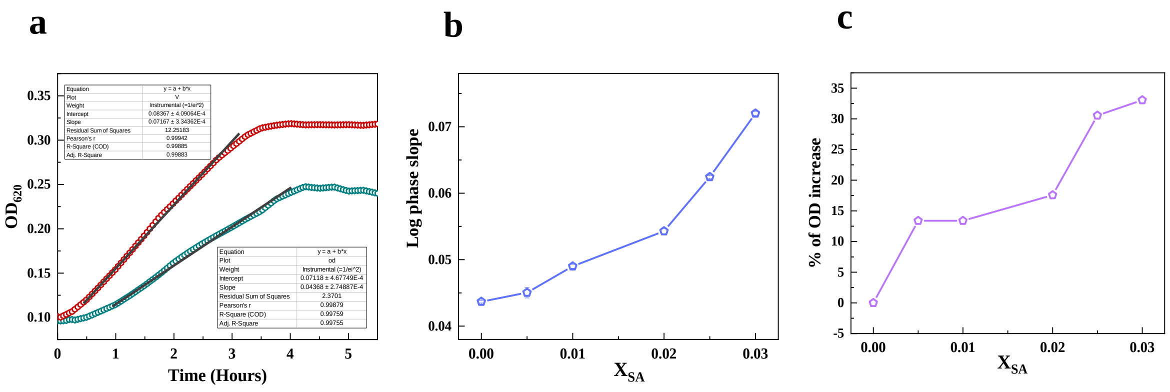


**Figure S2:** **(a)** Growth kinetics curves of bacterial cultures with and without SA incubation, showing a clear increase in the slope and hence a higher division rate after SA treatment. **(b)** Slopes extracted from the log phase for cultures incubated with different SA concentrations. **(c)** Percentage increase in OD at various SA concentrations, with a maximum change of approximately 33% observed at 0.03% SA.

all concentrations tested, as shown in Figure S2a. Figure S2b represents the slope dependence as a function of concentration of SA. The slope increased from 0.043 to 0.072, representing an approximate 67% enhancement, indicating a substantial increase in the specific growth rate of bacteria in the presence of SA. The slopes obtained at each concentration were further plotted as a function of the SA concentration, revealing a clear monotonic increase. Due to solubility limitations of SA, experiments could not be conducted at concentrations higher than 0.03%. The optical density at the stationary phase, which reflects the maximum cell density (carrying capacity), was also evaluated (Figure S2c). For *S. epi*, an increase of approximately 33% in OD at the stationary phase was observed at the highest SA concentration tested, suggesting that SA not only accelerates the growth rate but also supports a higher final bacterial population.

#### Derivation of the Gompertz equation for Bacterial growth kinetics

The Gompertz model is commonly used to describe bacterial growth kinetics.[1] The governing differential equation can be written as

$$\frac{1}{y}\frac{dy}{dt} = c\ln\left(\frac{a}{y}\right), \tag{S1}$$

where $y(t)$ represents the optical density at time $t$, $y_0$ is the initial value at $t = 0$, $a$ is the asymptotic steady-state value, and $r$ is the growth rate constant.

Rewriting Eq. S1 in terms of $\ln y$ and solving the resulting differential equation with the initial condition $y(0) = y_0$, the analytical solution can be expressed as

$$y(t) = a\exp\left[-\exp(-ct)\ln\left(\frac{a}{y_0}\right)\right]. \tag{S2}$$

For fitting experimental growth curves, it is convenient to express the solution in the following form

$$y(t) = y_0 + a\exp\left[-\exp(b - ct)\right], \tag{S3}$$

Here $a$ represents the asymptote at long time, corresponding to the maximum value approached by the growth curve. The parameter $b$ is related to the initial and asymptotic values through $e^b = \ln\left(\frac{a}{y_0}\right)$ and is therefore connected to $a$. The parameter $c$ is directly related to the growth rate and controls how rapidly the system approaches the asymptotic value.

Several parameters can be derived from the Gompertz model.[2,3] While some of these parameters are presented in the main manuscript, the remaining ones are described here and summarised in the Table S2 to provide a more complete picture of the various parameters. The time corresponding to the point of maximal growth rate, or inflection time ($T_{\text{inf}}$), was calculated from

$$T_{\text{inf}} = \frac{b}{c}, \tag{S4}$$

with the corresponding OD at this point given by

$$OD_{\text{inf}} = \frac{a}{e}. \tag{S5}$$

To further characterize growth, we determined the generation time ($T_g$), i.e. the time required for the bacterial population to double during exponential growth, using

$$T_g = \frac{\log(2)}{\mu_{\max}}. \tag{S6}$$

**Table S1:** Coefficients of the modified Gompertz model fitted to the growth curves of *S. epi* at different concentrations of SA.

| SA (%) | $y_0$ | $a$ | $b$ | $c$ |
|---|---|---|---|---|
| 0 | 0.096 ± 0.0003 | 0.113 ± 0.0004 | 1.92 ± 0.07 | 1.31 ± 0.05 |
| 0.005 | 0.092 ± 0.0009 | 0.149 ± 0.001 | 1.90 ± 0.07 | 1.17 ± 0.03 |
| 0.01 | 0.098 ± 0.001 | 0.157 ± 0.001 | 1.82 ± 0.38 | 1.20 ± 0.22 |
| 0.02 | 0.096 ± 0.001 | 0.175 ± 0.0003 | 1.76 ± 0.13 | 1.15 ± 0.05 |
| 0.025 | 0.103 ± 0.001 | 0.221 ± 0.001 | 1.35 ± 0.02 | 0.98 ± 0.01 |
| 0.03 | 0.098 ± 0.002 | 0.230 ± 0.002 | 1.31 ± 0.03 | 0.97 ± 0.01 |

**Table S2:** Growth parameters of *S. epi* derived from the modified Gompertz model at different concentrations of SA.

| SA (%) | $\mu_{\max}$ (1/h) | $\lambda$ (h) | $T_{\text{exp}}$ (h) | $T_{\text{inf}}$ (h) | $OD_{\text{inf}}$ | $T_g$ (h) |
|---|---|---|---|---|---|---|
| 0 | 0.054 ± 0.001 | 0.70 ± 0.05 | 2.07 ± 0.04 | 1.46 ± 0.05 | 0.04 ± 0.001 | 5.52 ± 0.08 |
| 0.005 | 0.064 ± 0.001 | 0.77 ± 0.05 | 2.32 ± 0.04 | 1.62 ± 0.05 | 0.05 ± 0.001 | 4.68 ± 0.06 |
| 0.01 | 0.069 ± 0.001 | 0.68 ± 0.07 | 2.27 ± 0.04 | 1.51 ± 0.27 | 0.058 ± 0.001 | 4.34 ± 0.31 |
| 0.02 | 0.074 ± 0.001 | 0.66 ± 0.08 | 2.36 ± 0.05 | 1.53 ± 0.08 | 0.064 ± 0.001 | 4.06 ± 0.07 |
| 0.025 | 0.079 ± 0.001 | 0.36 ± 0.01 | 2.77 ± 0.03 | 1.38 ± 0.01 | 0.08 ± 0.001 | 3.79 ± 0.02 |
| 0.03 | 0.082 ± 0.001 | 0.32 ± 0.01 | 2.80 ± 0.05 | 1.36 ± 0.01 | 0.085 ± 0.001 | 3.66 ± 0.03 |

To assess whether SA exerts a similar effect on other bacteria, we examined growth kinetics at the same concentration in *E. coli*, a Gram-negative pathogenic bacterium. As shown in Figure S3, the effects were minimal compared with those observed in *S. epi*. As seen in Figure S3, there is no noticeable change in slope during the log phase and only minimal variation in OD at the highest concentration (0.03 %), suggesting that the effects of SA on *E. coli* are not very significant.

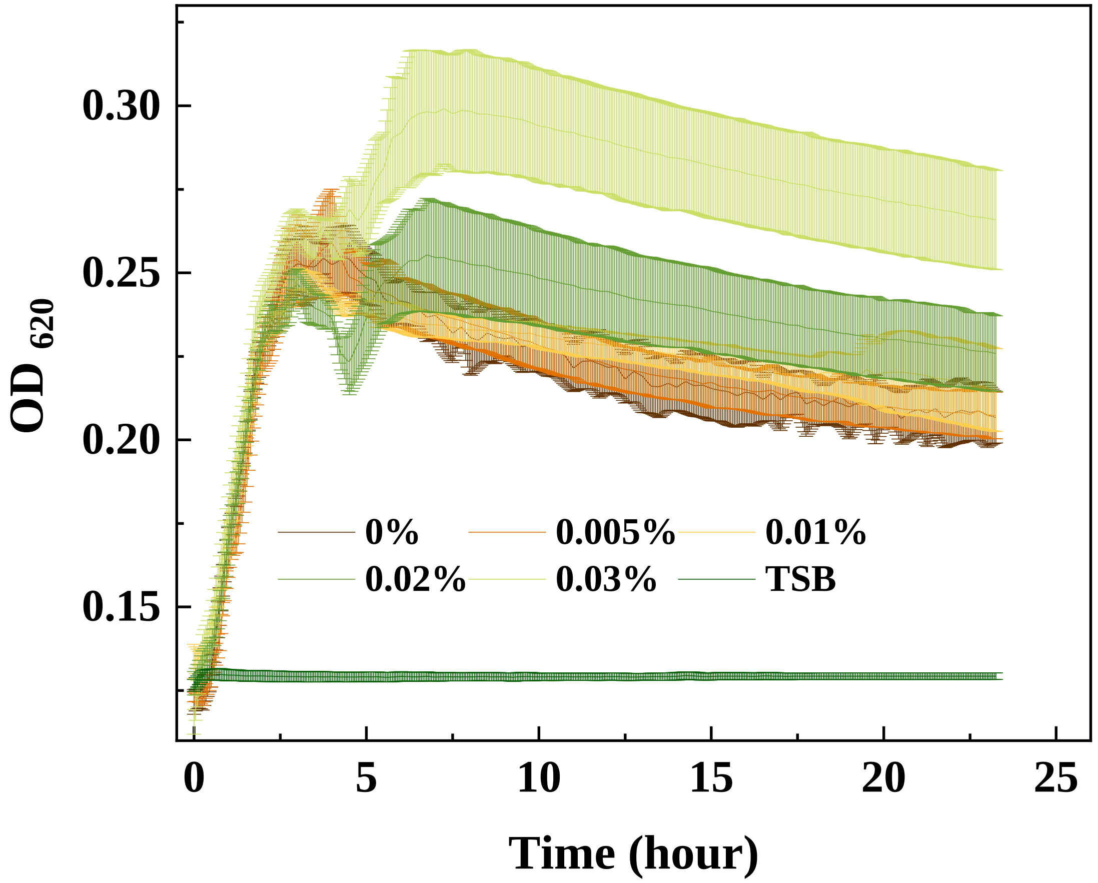


**Figure S3:** Growth kinetics curves of bacterial cultures with and without SA incubation, showing a marginal increase in the growth kinetics curves after SA treatment, showing the effect on *E.coli* is not as effective as *S. epi.*

### Fluorescence Correlation Spectroscopy (FCS)

The average particle number in the observation spot is denoted by <N>and the fluctuation in particle number is represented by ΔN(t). The total number of particles at any given time can be written as:

$$N(t) =< N > +\Delta N(t) \tag{S7}$$

By measuring the intensity of the emission (I (t) =<I>+ ΔI(t)); where <I>and Δ I (t) represent the intensity and fluctuation over time, respectively, we can calculate the fluctuations of the number of fluorescent molecules entering and leaving the focal plane of a confocal microscope observation spot during excitation. In FCS, the autocorrelation function G($\tau$) from the intensity signal I (t) measured under the microscope is calculated using;[4]

$$g_2(\tau) = \frac{\langle \Delta I(t) \Delta I(t+\tau) \rangle_t}{\langle I(t) \rangle_t^2} \tag{S8}$$

where $<>_t$, denotes that the physical quantity was averaged over the time variable t. The correlation data were normalized and fitted to a single or multi-component 2D diffusion equation, the following equation:

$$g_2(\tau) = G_\infty + \left[ \frac{1}{N} \frac{\left(1 - \theta_t + \theta_t e^{-\frac{t}{\tau_t}}\right)}{1 - \theta_t} \frac{\rho}{\left(1 + \frac{t}{\tau_D}\right)^\alpha} \right] \tag{S9}$$

Here $G_\infty$ is the background of the curve where the autocorrelation decays to 0. $\alpha$ is the anomaly parameter, which is very close to 1 for Brownian diffusion and deviates from 1 in the case of the sub-diffusive or super-diffusive nature of diffusion. $\theta_t$ is the fraction of molecules that are present in the triplet state, $\tau_t$ is the relaxation time in the triplet state, $\rho$ is the fraction of diffusing molecules, which is unity in these fits, and $\tau_D$ is the diffusion time. The Quickfit software was used to extract the $\tau_D$ parameter from Equation S9. The anomaly parameter $\alpha$ values were approximately $0.9 \pm 0.1$. Once the $\tau_D$ values are known, the diffusivity $D$ can be evaluated using the following equation

$$D = \frac{d^2}{\tau_D 8 \ln 2} \tag{S10}$$

where $d$ represents the full width at half maximum (FWHM) of the Gaussian beam. $\tau_D$ is the diffusion time, and D is the diffusion coefficient of the Nile Red molecules in the lipid environment.[5] The model autocorrelation profiles with fits are given in Figure S4b.

Fluorescence correlation spectroscopy (FCS) was performed on bacteria sampled at different growth phases for pristine and SA-incubated cells. Measurements were taken during the lag phase (0–2 hours), the late log phase (approximately 4 hours, corresponding to the onset of saturation), and the stationary phase (8–10 hours), allowing for a comparison of the diffusion behaviour across the growth cycle. As shown in Figure S4a, pristine bacteria exhibit a slightly reduced diffusion coefficient in the stationary phase relative to the log and late-log phases. This decrease may be linked to the enhanced production of cardiolipin, a phospholipid with four acyl chains that promotes the tighter packing of membrane lipids, thereby increasing membrane rigidity.[6,7]

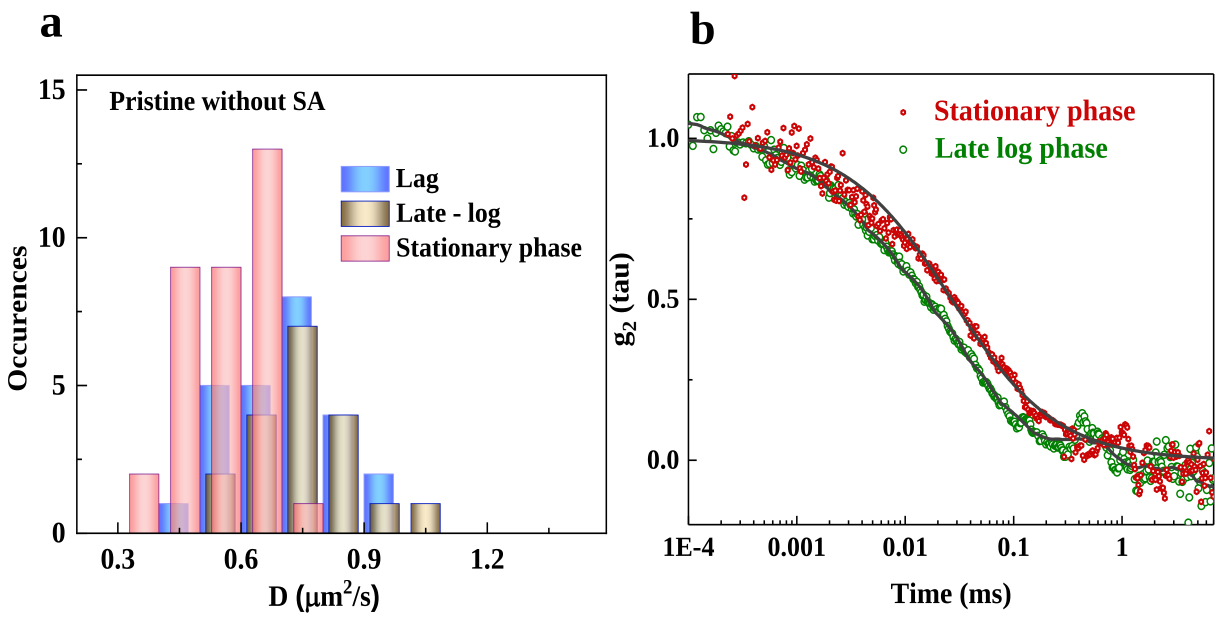


**Figure S4: (a)** Histogram of diffusion coefficients for pristine bacteria showing a noticeable shift toward lower values in the stationary phase compared to the late-log phase. **(b)** The corresponding autocorrelation curves for pristine cells in the late-log and stationary phases indicate slower dynamics in the stationary phase, consistent with reduced membrane fluidity.

We then collected growth profiles for bacteria incubated with SA during both the late log phase and the stationary phase to determine whether they exhibited behaviour comparable to that of pristine cells. Similar trends were observed following SA incubation, indicating that the overall pattern of changes in the diffusion coefficient across growth phases is preserved. Quantitatively, the extent of fluidization approached approximately 89% in both phases, as shown in Figure S5, suggesting that SA exerts a consistent influence on membrane dynamics across these stages.

#### Agarose pad experiments

The diffusion coefficient is an important parameter for assessing changes in membrane dynamics. It can also serve as an indicator of whether bacteria revert to their native state once

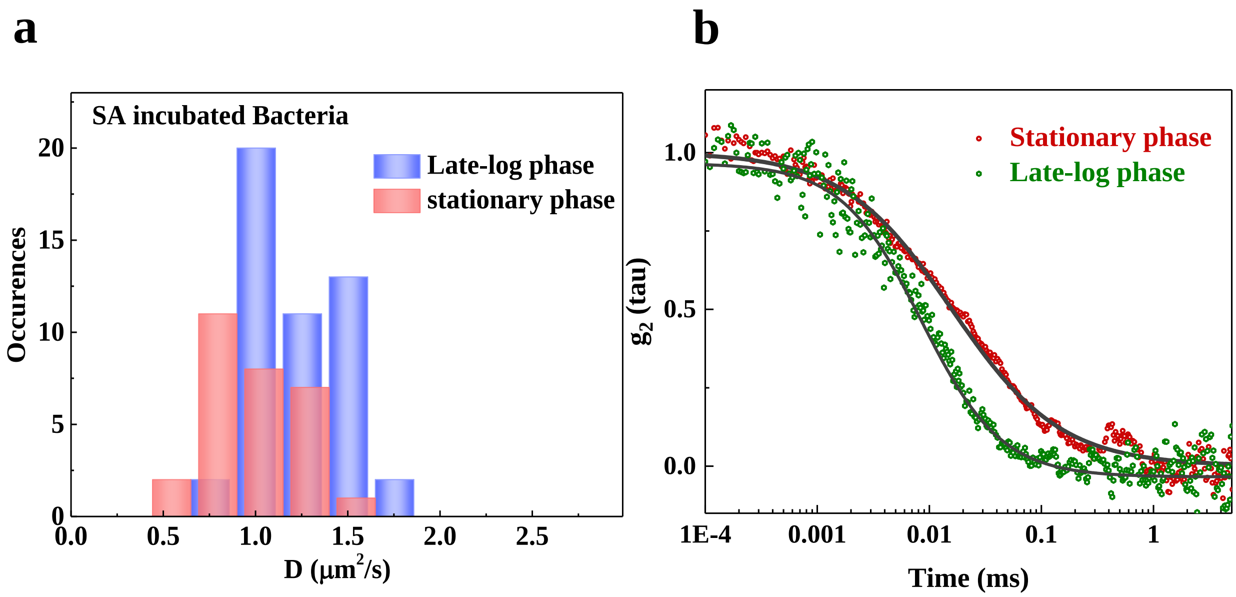


**Figure S5:** **(a)** Histogram of diffusion coefficients for SA-treated cells, showing a shift toward lower values in the stationary phase compared to the late-log phase. **(b)** Corresponding autocorrelation curves indicate slower dynamics in the stationary phase, consistent with reduced membrane fluidity even after incubation with SA.

the SA effect is removed, as well as how long the effect of SA persists, based on differences in the diffusion coefficient between pristine and SA-incubated cells. To evaluate this, bacteria were incubated with SA and then spread on an agarose pad without SA. Diffusion coefficients were measured for four successive generations on the agarose pad. Since SA could not be mixed with agarose due to solubility issues, this setup effectively served as an in-vitro assay to test how long the effect of SA is retained by the bacteria. As shown in Figure S6a, four generations of *S. epi* were tracked, and the dye concentration was found to be sufficient to perform FCS experiments. FCS measurements were carried out across generations, and the diffusion coefficient values are plotted in Figure S6b, which shows no noticeable change. To further examine how many generations this effect of SA persists, bacteria were cultured in liquid medium and diffusion coefficients were calculated for the 1st and 12th generations, as shown in Figure S6c. Interestingly, the diffusion coefficient values reverted to those observed in bacteria incubated without SA (**??**a).

### Viscosity calibration

Viscosities were determined using a previously reported viscosity–lifetime calibration equation.[8] To calibrate the constants, fluorescence lifetimes were measured from decay curves obtained in a series of glycerol/methanol mixtures with known viscosities. The calibration curve for different concentrations of glycerol/methanol mixtures is shown in Figure S7. Fluorescence lifetime values obtained from these mixtures were plotted against their known viscosities, and the data were fitted to a straight line. From this plot, the slope and intercept were extracted and applied to the Förster–Hoffmann (FH) equation to determine the viscosity of unknown samples. In our case, the slope and intercept values derived from the calibration curve were substituted into the equation to calculate the membrane viscosity of bacteria labelled with and without SA treatment. From the calibration curve (see Figure S7), the extracted slope and intercept were converted into the constants of the FH equation, yielding values of 0.489 and 0.456, respectively. Substituting these constants into the relation gives the following equation,

$$\log \eta = \frac{\log \tau_1 + 0.489}{0.456} \tag{S11}$$

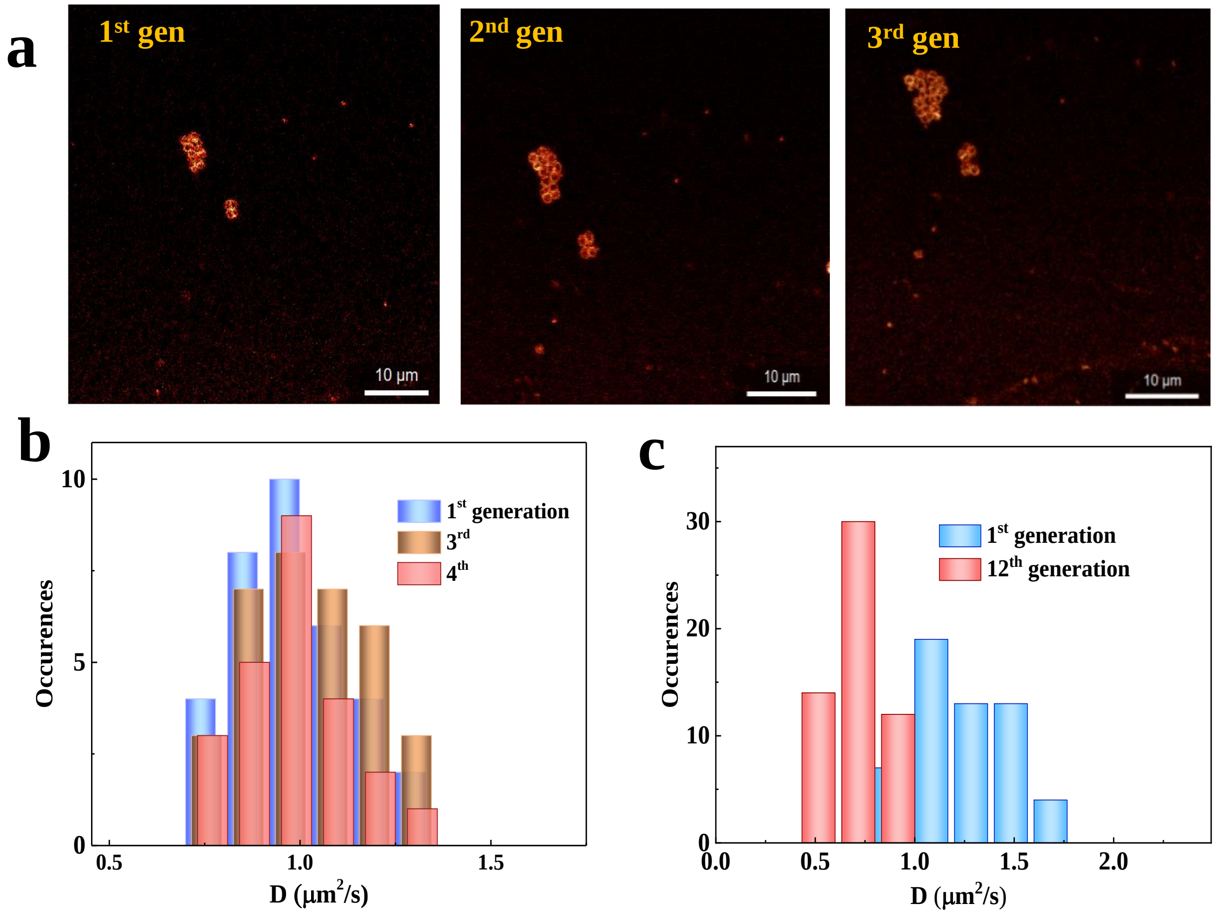


**Figure S6:** **(a)** Confocal images of bacterial colonies grown on agarose pads across successive generations (1st, 2nd, and 3rd), with the cells labelled with NR to visualize the membrane**(b)** Histograms of diffusion coefficients indicate no significant changes over four generations when experiments are performed on agarose pads. **(c)** A significant reversal of the diffusion coefficient toward the pristine state is observed, suggesting that the cells attempt to revert back when the experiments are extended to 12 generations in TSB medium. The number of profiles obtained was higher for bacteria incubated in liquid medium, whereas experiments performed on agarose pads yielded fewer profiles due to cell division limiting the number of stable measurement regions.

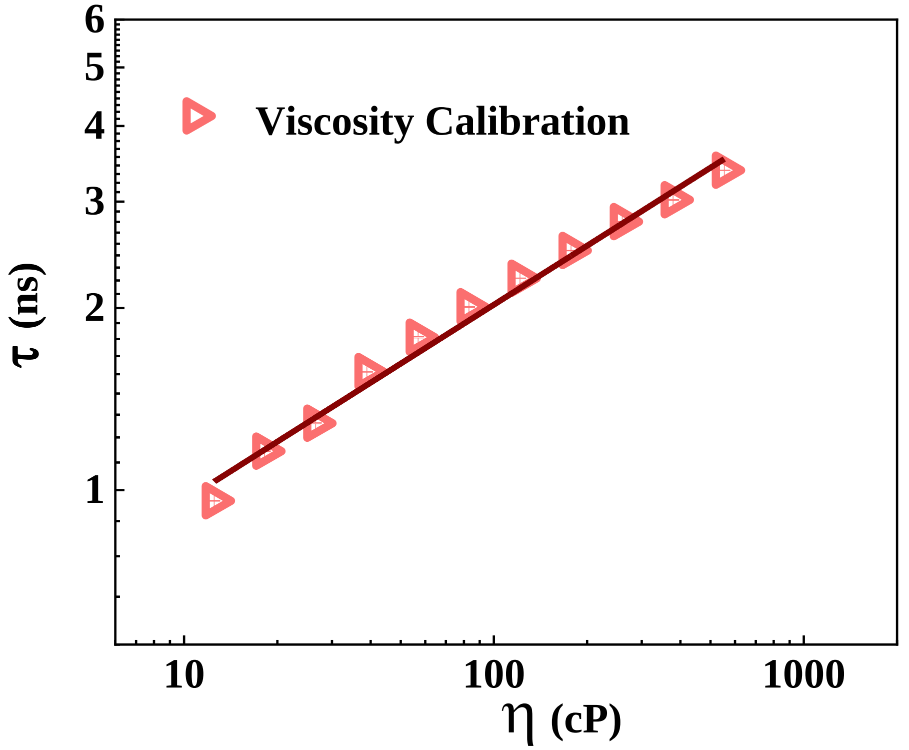


**Figure S7:** Log–log plot of fluorescence lifetime ($\tau$) of BODIPY-C12 versus viscosity ($\eta$) for methanol–glycerol mixtures. The linear fit follows the FH relation and provides calibration constants for converting lifetime data to membrane viscosity.

#### Lifetime measurements

Time-resolved spectroscopy is used to investigate dynamic processes in materials and chemical compounds by monitoring how their spectroscopic properties evolve over time. By illuminating a sample with pulsed lasers, it becomes possible to observe phenomena that occur on extremely short time scales, even down to the femtosecond range. The lifetime of an excited molecule refers to the duration it takes for a group of such molecules to decay to $1/e$ of their initial excited state population. Consequently, the fluorescence lifetime denotes

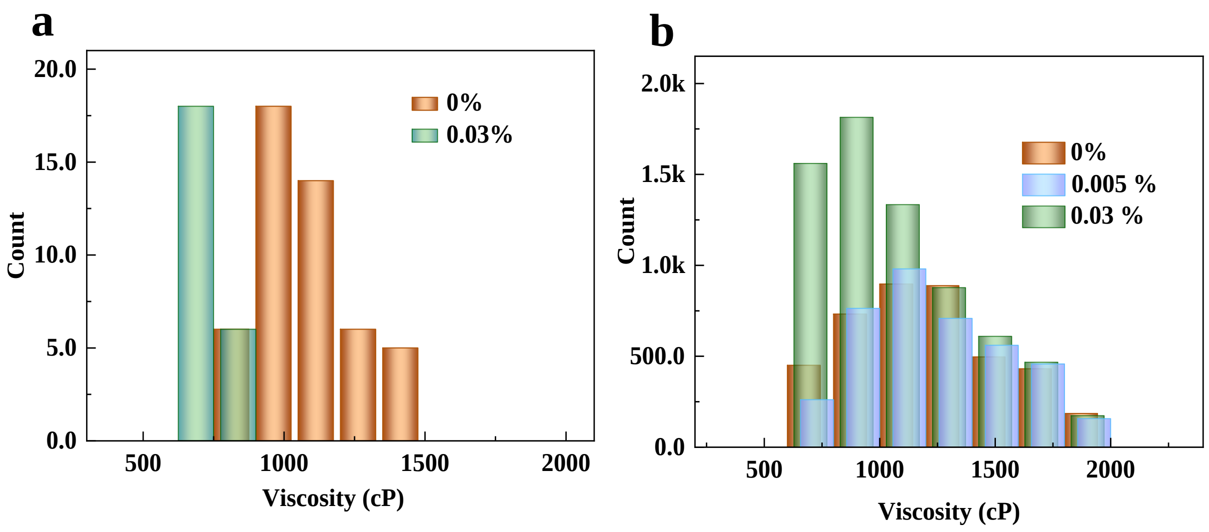


**Figure S8:** **(a)** Histogram of membrane viscosity directly measured using the BODIPY C12 molecular rotor in FLIM. **(b)** Histogram of viscosity calculated from FCS diffusion data using the Saffman-Delbrück (SD) model. Both methods independently show a significant decrease in membrane viscosity upon SA treatment.

the average time a molecule remains in its excited state before emitting a photon. The decay of fluorescence is commonly represented by a first-order rate equation, as shown in Equation S12.

$$A_t = A_0 e^{-\frac{t}{\tau_t}} \tag{S12}$$

Here, $A_t$ is the initial intensity of fluorescence, which decays with time t, and $\tau_t$ is the lifetime of the fluorescence. Fitting was performed in the FLIM software integrated with the Picoquant system, and fluorescence lifetime images were displayed in pseudo-colour (main manuscript). The goodness of fit parameter was obtained with a precision of $\chi^2 \approx 0.9981$. To investigate the lifetime (viscosity) trends in detail and to get a good $\chi^2$ for fitting, biexponential decays were used to get good fits.

To compare our diffusion coefficient obtained using FCS methods to the membrane viscosity measured, the Saffman–Delbrück (SD) model[9] was used. This model uses a fluorescent probe as a cylindrical object diffusing within the fluid-like cell membrane. This model relates the diffusion coefficient $D$ to the membrane's two-dimensional viscosity $\mu_m$ through the following equation

$$D = \frac{k_B T}{4\pi \mu_m h}\left[\ln\left(\frac{\mu_m}{\eta_s a}\right) - \gamma\right], \tag{S13}$$

Here $k_B$ is Boltzmann's constant ($\approx 1.38 \times 10^{-23}$ J/K), T is absolute temperature (here, 293 K), $\eta_s$ is the viscosity of the surrounding bulk fluid, r is the effective hydrodynamic radius of the diffusing particle ($\sim$ 0.6 nm[10]), $\gamma \sim 0.5772$ is the Euler–Mascheroni constant.

The two-dimensional membrane viscosity $\mu_m$ is related to the three-dimensional (shear) viscosity $\eta_m$ by:

$$\mu_m = \eta_m\, h \tag{S14}$$

where h is the membrane thickness (typically 3.5-4 nm for bacterial membranes). By calculating the diffusion coefficient from our FCS measurements and applying the SD model, we were able to estimate the corresponding membrane viscosity.

This approach provides an independent means of validating the viscosity values obtained from our direct measurements. From Figure S8, which shows the viscosity values converted from the diffusion coefficients using the SD equation, it can be seen that for the pristine samples, the viscosity values are in close agreement. After incubation with SA, the qualitative trend remains the same; however, the viscosity values derived from the diffusion coefficients show larger changes, possibly because of local effects and the varying positions of the dye molecules within the lipid molecules.

### Probing Peptidoglycan Dynamics Using FLIM

For labelling the peptidoglycan, wheat germ agglutinin (WGA) conjugated with Alexa Fluor 647 was used,[11] with the emission detector set to record in the 650–700 nm range. For the labelling process, 0.5 OD bacteria were incubated with the WGA Alexa flour 647 for 1 hour at 37°C with shaking. Following incubation, the bacteria were washed multiple times with PBS to remove any unbound dye. The labelled bacteria were then spread onto a PLL-coated glass substrate, washed again with PBS to ensure minimal background fluorescence, and then imaged under the microscope. Fluorescence lifetime measurements were recorded both in the presence and absence of SA incubation to assess any changes. Since WGA does not exhibit viscosity dependence due to its molecular structure, the FH equation could not be applied to determine the constants required to extract viscosity values from the measured fluorescence lifetimes ($\tau$). Therefore, the lifetime values are plotted directly in order to assess any changes arising from alterations in the microenvironment caused by incubation with SA. This approach allows us to visualize and interpret how the local surroundings of the probe are affected, providing insight into possible interactions or structural modifications induced by SA. As can be clearly seen from the images in Figure S9a and b, as well as from the lifetime decay curve in panel c, which provides a quantitative representation of the observations in panels a and b of Figure S9, no significant alterations are observed in the PG layer.

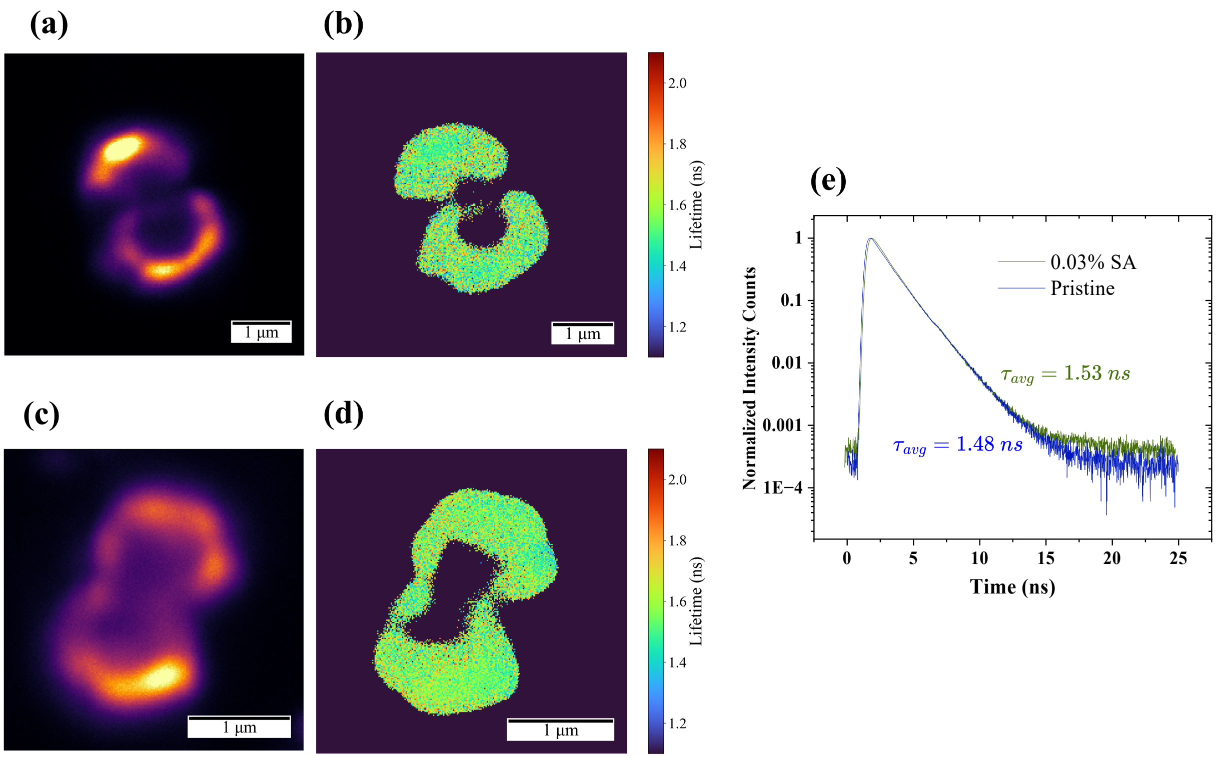


**Figure S9:** (a & b) Intensity and lifetime images for pristine *S. epi* labelled with WGA, a peptidoglycan binding dye.(c & d) *S. epi* incubated with 0.03% of SA, where the maximum changes were observed.(e) Average lifetime extracted by selecting various regions of interest on the bacteria for pristine *S. epi*.(f) With 0.03% of SA incubation. (g) Decay profiles for pristine *S. epi* and for the bacteria incubated with 0.03% of SA, showing significant variation. The lifetime changes observed are opposite to the trend seen in the case of membrane lifetime, labelled using a membrane-labelling dye.

### Atomic Force Microscopy

In the main manuscript, we presented AFM data for bacteria collected on glass substrates coated with Vectabond. To evaluate the reproducibility of the data and to achieve better bacterial adhesion, we repeated the experiments using high-density PLL (Sigma) as the coating material. As shown in S10b and c, an increase in Young's modulus, calculated using Hertz-model,[12] is observed, indicating membrane stiffening upon incubation with SA.[13,14]

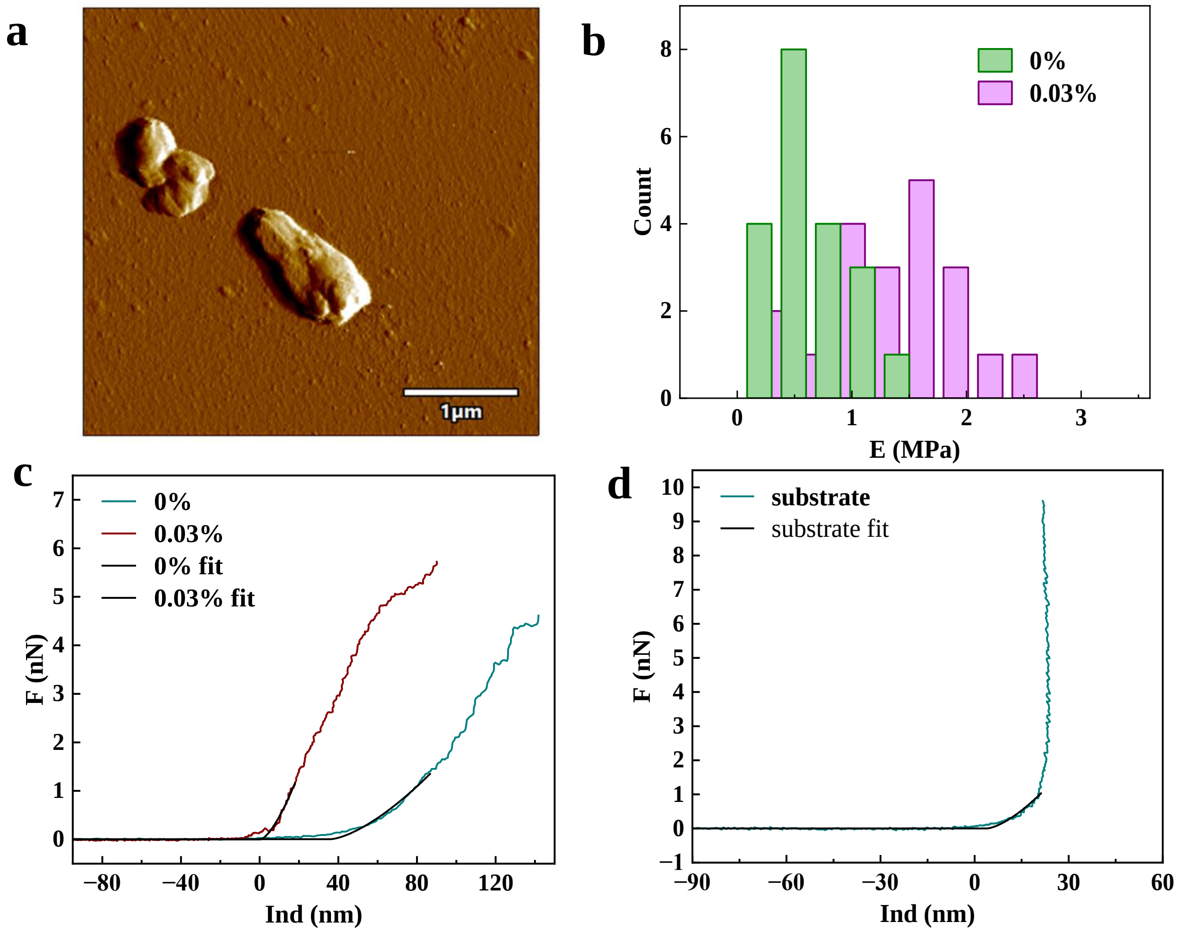


**Figure S10:** **(a)** Amplitude image of *S. epi* recorded under liquid conditions on a high-density PLL-coated substrate. **(b)** Histogram of elastic modulus values for pristine and SA-treated bacteria, showing an increase in stiffness upon SA incubation. **(c)** Representative F-D curves for pristine and SA-treated cells with corresponding Hertz-model fits. **(d)** F-D curve obtained on the PLL-coated substrate with its fit.

#### Coarse-grained Martini Simulation Results

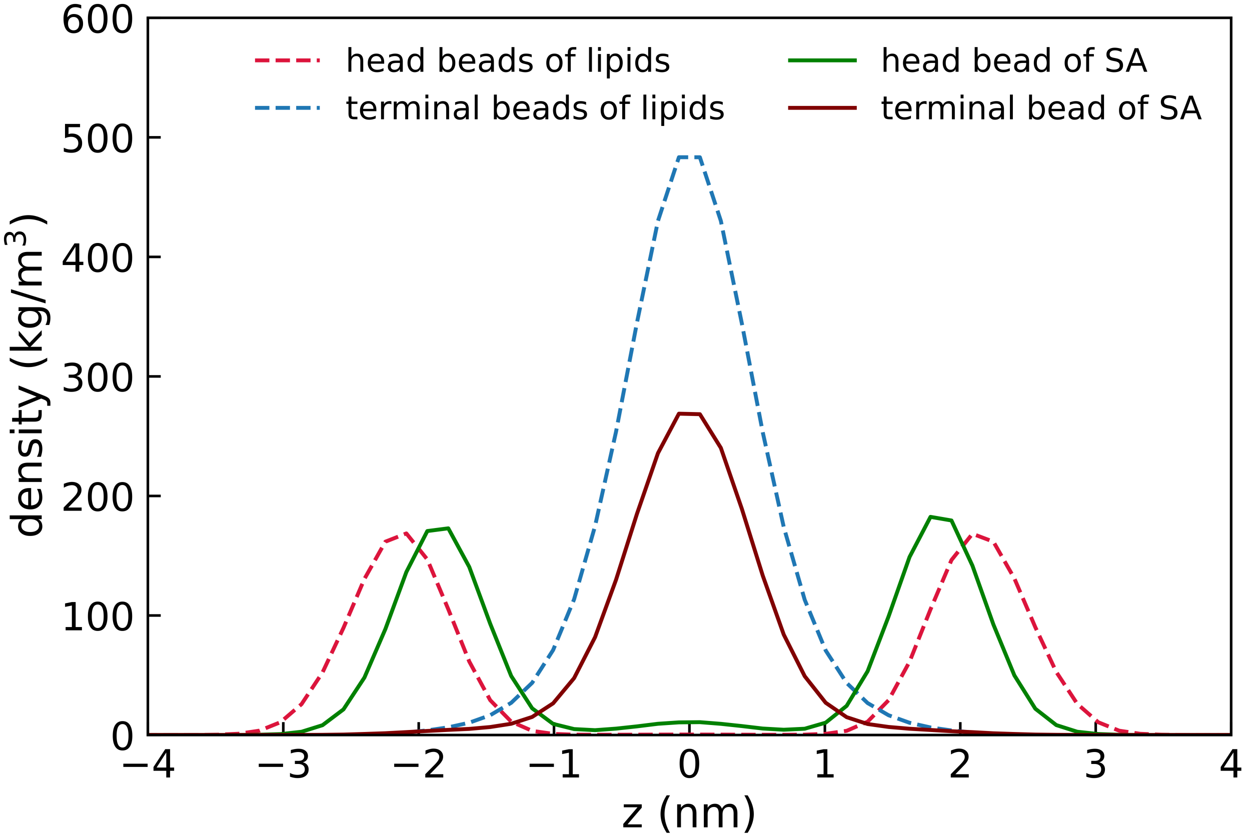


**Figure S11:** Profiles showing the densities of headgroups and terminal beads of lipids and SA molecules across the bilayer normal in simulation containing 30 molecules of SA. The density values for SA are scaled up by a factor of 20 for visualization purpose. The density of the head bead of SA in the core of the bilayer is due to flip-flop of SA molecules across the membrane leaflets.

The data for mean squared displacement are shown in Figure S12 for 10 ns long Martini simulations containing 2, 10 and 30 molecules of SA.

The lateral diffusion coefficients (D) for membrane lipids are computed by linear regression of MSD data obtained in Martini simulations. Table S3 below indicates the D values for simulation cases with 2, 10 and 30 SA molecules.

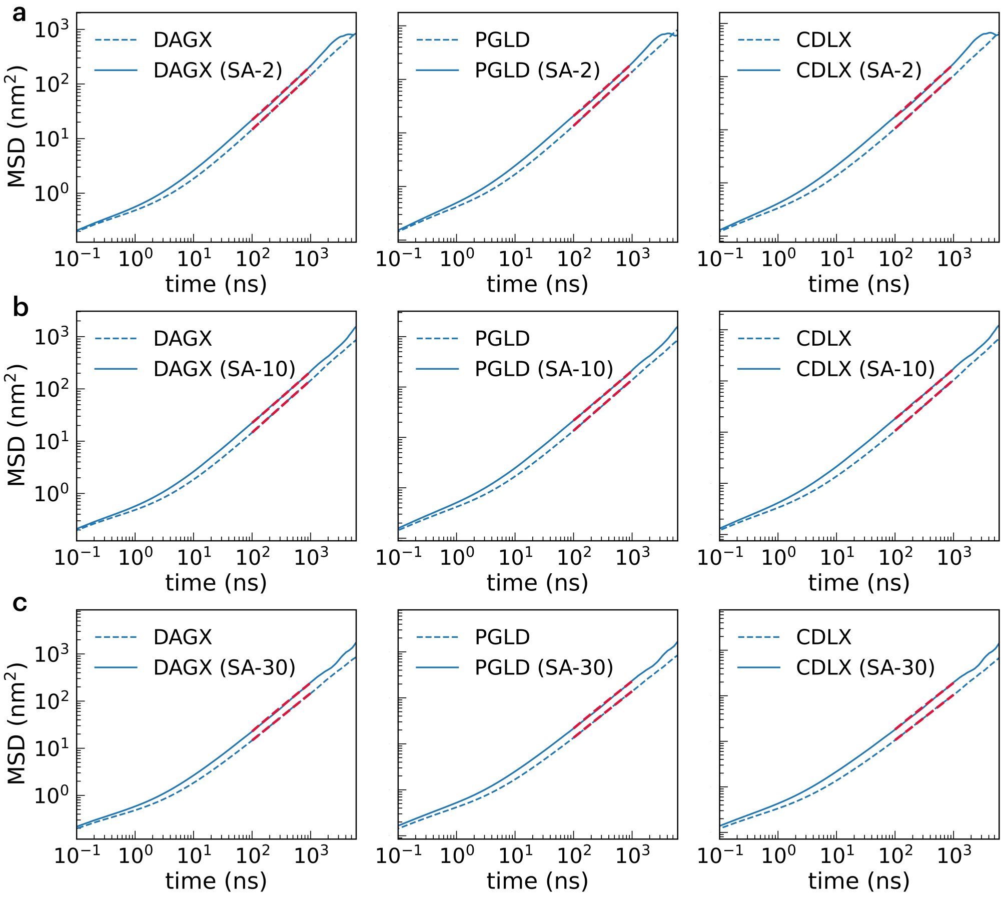


**Figure S12:** Mean squared displacements (MSD) for individual lipids - DAGX, PGLD and CDLX in Martini simulations with (a) 2 molecules of SA, (b) 10 molecules of SA, and (c) 30 molecules of SA. The blue dotted lines represent the MSD for the membrane lipids without SA. The fits for MSD $= 4Dt$ are indicated in red dotted lines.

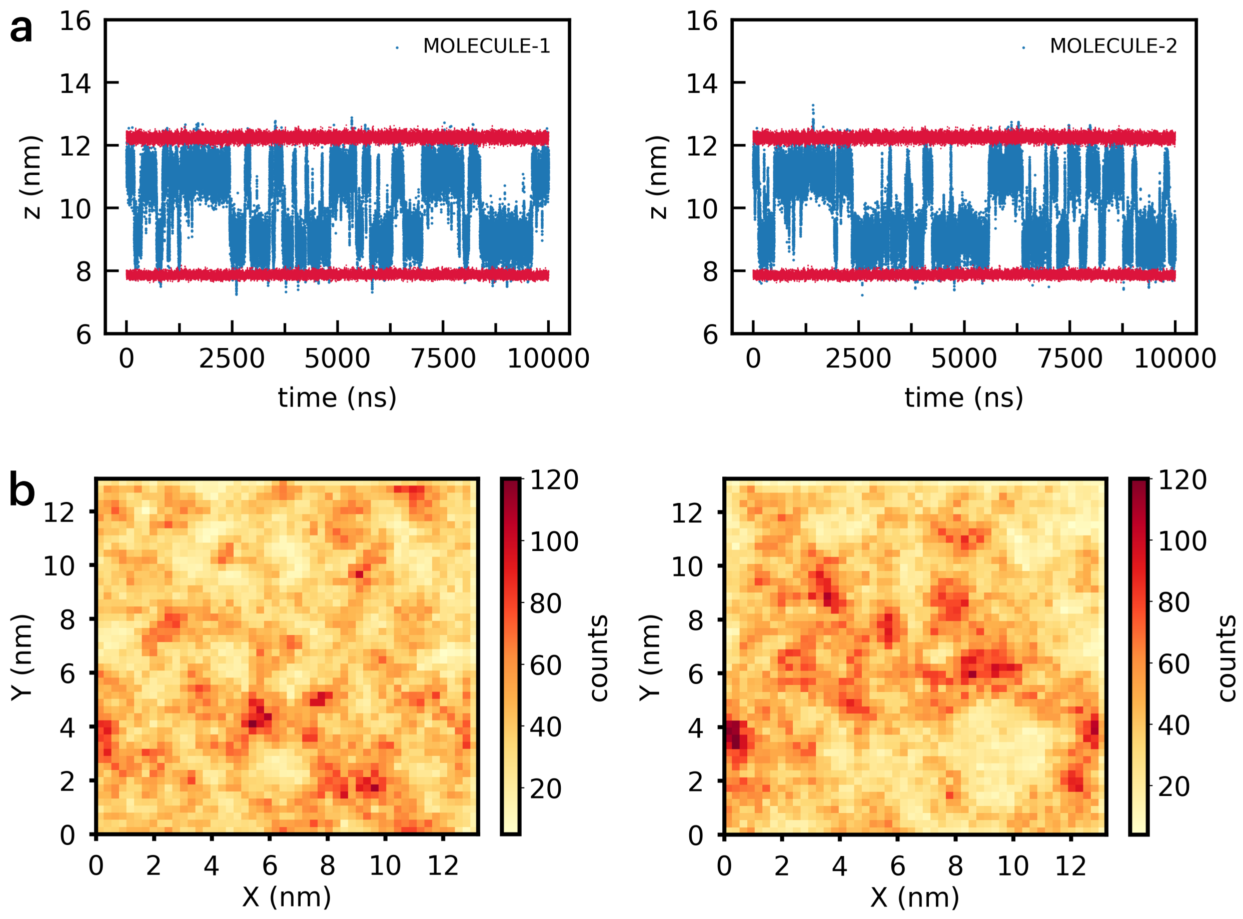


**Figure S13:** Center of mass motion for SA molecules in simulation with 2 molecules of SA. (a) z-component of the center of mass trajectory of SA molecules (blue) and lipid headgroup beads (red) indicating the lipid-water interfaces and the flip-flop events of SA across the bilayer leaflets within the core of the membrane. (b) Sampling of individual SA molecules in X-Y membrane plane for 10 $\mu$s long simulation run.

**Table S3:** Lateral diffusion coefficient extracted from $MSD = 4Dt$ fits for simulations of the bare membrane and membranes containing of 2 (SA-2), 10 (SA-10) and 30 (SA-30) molecules of stearic acid. The best fits with goodness $R^2 > 0.99$ were obtained, and the diffusion coefficient is reported in units of $10^{-7}$ $cm^2/s$. The last column reports the molar weighted average value of the diffusion coefficient averaged over the membrane lipids and the simulation runs.

| System | DAGX | PGLD | CDLX | SA | Average |
|---|---|---|---|---|---|
| **Bare membrane (run 1)** | 3.67 | 3.48 | 2.63 | – | $3.18 \pm 0.20$ |
| **Bare membrane (run 2)** | 3.25 | 3.03 | 2.39 | – | |
| **Membrane + SA-2 (run 1)** | 5.28 | 4.94 | 4.27 | 7.92 | $5.00 \pm 0.28$ |
| **Membrane + SA-2 (run 2)** | 5.63 | 5.51 | 4.65 | 9.19 | |
| **Membrane + SA-2 (run 3)** | 5.63 | 5.51 | 4.65 | 9.19 | |
| **Membrane + SA-10 (run 1)** | 5.34 | 5.21 | 4.25 | 8.02 | $5.26 \pm 0.20$ |
| **Membrane + SA-10 (run 2)** | 5.44 | 5.12 | 4.74 | 8.04 | |
| **Membrane + SA-10 (run 3)** | 5.78 | 5.61 | 4.91 | 9.60 | |
| **Membrane + SA-30 (run 1)** | 6.15 | 5.80 | 4.86 | 8.62 | $5.85 \pm 0.29$ |
| **Membrane + SA-30 (run 2)** | 6.60 | 6.25 | 5.53 | 8.66 | |
| **Membrane + SA-30 (run 3)** | 5.86 | 5.65 | 4.77 | 8.55 | |

### References


[1] Y. M. G. Montes, E. R. V. Calle, S. G. S. Terán, M. R. C. García, J. C. R. Nájera and M. R. L. Vera, *Journal of the Science of Food and Agriculture*, 2024, **104**, 1258–1270.

[2] J. Wang and X. Guo, *Biotechnology Advances*, 2024, **72**, 108335.

[3] T. Chatterjee, B. K. Chatterjee, D. Majumdar and P. Chakrabarti, *Biochimica et Biophysica Acta (BBA)-General Subjects*, 2015, **1850**, 299–306.

[4] P. Sharma, S. Parthasarathi, N. Patil, M. Waskar, J. S. Raut, M. Puranik, K. G. Ayappa and J. K. Basu, *Langmuir*, 2020, **36**, 8800–8814.

[5] G. Singh, A. C. Chamberlin, H. R. Zhekova, S. Y. Noskov and D. P. Tieleman, *Journal of Chemical Theory and Computation*, 2016, **12**, 364–371.

[6] S. Hiraoka, H. Matsuzaki and I. Shibuya, *FEBS letters*, 1993, **336**, 221–224.

[7] T. Koprivnjak, D. Zhang, C. Ernst, A. Peschel, W. Nauseef and J. Weiss, *Journal of bacteriology*, 2011, **193**, 4134–4142.

[8] J. T. Mika, A. J. Thompson, M. R. Dent, N. J. Brooks, J. Michiels, J. Hofkens and M. K. Kuimova, *Biophysical journal*, 2016, **111**, 1528–1540.

[9] P. Saffman and M. Delbrück, *Proceedings of the National Academy of Sciences*, 1975, **72**, 3111–3113.

[10] N. C. for Biotechnology Information, *PubChem Compound Summary for CID 65182, Nile Red*, 2025, Retrieved August 27, 2025 from `https://pubchem.ncbi.nlm.nih.gov/compound/Nile-Red`.

[11] W. Vollmer, D. Blanot and M. A. De Pedro, *FEMS microbiology reviews*, 2008, **32**, 149–167.

[12] E. K. Dimitriadis, F. Horkay, J. Maresca, B. Kachar and R. S. Chadwick, *Biophysical journal*, 2002, **82**, 2798–2810.

[13] R. G. Bailey, R. D. Turner, N. Mullin, N. Clarke, S. J. Foster and J. K. Hobbs, *Biophysical journal*, 2014, **107**, 2538–2545.

[14] R. Han, W. Vollmer, J. D. Perry, P. Stoodley and J. Chen, *Nanoscale*, 2022, **14**, 12060–12068.
